# Deep Receptor Scanning Reveals General Sequence Constraints on GPCR Biosynthesis

**DOI:** 10.1101/2025.09.19.677468

**Authors:** Austin Tedman, Muskan Goel, Sohan Shah, Mathew K. Howard, Laura M. Chamness, Antonio Bonifasi, Ismalia Adams, Jacklyn M. Gallagher, Wesley D. Penn, Katarina Nemec, Eli F. McDonald, Brianna N. Corman, J. Paul Robinson, Carol Beth Post, Patricia L. Clark, M. Madan Babu, Aashish Manglik, Charles P. Kuntz, Willow Coyote-Maestas, Jonathan P. Schlebach

## Abstract

G protein-coupled receptors (GPCRs) mediate a variety of signaling pathways and are the most common pharmacological targets. While advances in structural biochemistry have provided deep functional insights into key receptors, many of the 800+ human GPCRs remain understudied. We introduce a versatile “deep receptor scanning” platform that can be used to experimentally characterize 766 human GPCRs and 174 known GPCR splice variants in parallel. We use this platform to quantitatively characterize the relative abundance of canonical and alternative receptor transcripts, their translational efficiency, and the plasma membrane expression of each receptor in the context of a recombinant pool of HEK293T cells expressing individual GPCRs. We then employ machine learning to identify specific structural features that modulate GPCR expression. This experimental platform and informatic approach are compatible with a variety of assays and can be used to efficiently explore the biochemical and pharmacological properties of the GPCRome.

## Introduction

G Protein-Coupled Receptors (GPCRs) constitute the largest family of human genes and modulate a variety of intracellular signaling cascades in response to extracellular stimuli. The disruption of these signaling pathways is tied to an extensive list of pathological conditions. For this reason, GPCRs are targeted by approximately 35% of current therapeutics.^1,2^ Advances in the cellular biology and structural biochemistry of GPCR signaling over the past few decades have revealed how small molecules tune their interactions with transducers in a manner that modulates downstream signaling cascades.^3–5^ While numerous generalizable structure-function relationships have emerged, relatively little is known about how variations in the sequence, structure, and splicing of GPCRs impact their expression, localization, and/ or turnover.

Efforts to dissect the molecular determinants of GPCR specificity have benefitted tremendously from advances in cellular biochemistry,^6^ structural biology, dynamic measurements/ simulations,^7–10^ and molecular modeling.^11^ Modern breakthroughs in crystallography^12,13^ and electron microscopy^14,15^ have facilitated structure determination for hundreds of GPCRs and GPCR signaling complexes. Additionally, molecular dynamic simulations have revealed how small molecules prompt receptors to transition from one conformation to another. Experimental structures have also empowered comparative modeling and generative AI approaches, which can now be used to generate realistic structural models of most receptors and many important signaling complexes. Importantly, the accuracy of these models has reached a level that is now sufficient for computational drug discovery. Moreover, cryo-EM and structural modeling are beginning to reveal how intracellular interfaces encode specificity for certain transducers.^11^ Nevertheless, the prevailing paradigms in receptor biology are built upon deep mechanistic studies of a limited number of well-studied GPCRs. New methodological approaches to explore the GPCRome could provide new opportunities to evaluate which of these paradigms are fully generalizable across diverse receptor classes.

While emerging machine learning approaches could conceivably address several eminent challenges in GPCR biology, their ultimate capabilities are likely to be constrained by the scope of the available training data (or lack thereof). And while there are rich data that describe the relationships between the sequence, structure, and biological function of many receptors, most GPCRs lack any sort of standardized experimental descriptors of their expression, trafficking, and/ or signaling activity in the cell. These understudied aspects of GPCR biology can have a profound impact on the evolutionary fitness of these receptors. Nevertheless, current high throughput GPCR platforms have been optimized for the measurement of specific biochemical or cellular readouts.^16,17^ For instance, brute-force approaches such as the PRESTO-Tango platform were designed to survey the modulation of several hundred receptors to small molecules of interest.^17^ While immensely useful, this arrayed receptor platform requires specialized robotic resources to scale and can only be used to detect activation events that recruit a chimeric β-arrestin transducer. Sequencing-based transcription readouts can be used to track the parallel activation of many receptors in a pooled experimental format,^18–20^ though platforms such as PRESTO-Salsa first require the production of hundreds of individual stable cell lines. Efforts to generate comprehensive experimental descriptors that span the GPCRome will therefore require more versatile high-throughput approaches.

In the following study, we describe a versatile experimental platform that leverages emerging genetic technologies from deep mutational scanning to characterize up to 766 canonical human GPCRs and 174 experimentally validated GPCR splice variants in parallel. Using this “deep receptor scanning” (DRS) approach, we characterize the transcript abundance, translational dynamics, and plasma membrane expression levels of 940 unique receptors and receptor isoforms in HEK293T cells. We identify receptors that are encoded by unstable transcripts and identify systematic differences in the translation dynamics of distinct classes of GPCRs. Our deep receptor scanning data reveal that the plasma membrane expression of these receptors varies by four orders of magnitude. Using various machine learning approaches, we comprehensively identify the general structural features that limit plasma membrane expression across the GPCR family. This experimental characterization of the GPCRome reveals generalizable insights into GPCR biosynthesis while establishing a versatile high-throughput platform that can be used to characterize numerous other aspects of GPCR activity at scale.

## Results

### Curation and Design of an Open-Source GPCR Library

The applications of current experimental GPCRome platforms are constrained by both the design of genetic libraries and the requisite expression systems. One leading platform, PRESTO-Tango, consists of a plasmid library encoding 315 non-olfactory receptors. However, this collection features codon-optimized versions of each open reading fame (ORF) that are fused to an exogenous signal peptide, an N-terminal epitope tag, and a lengthy C-terminal domain that recruits arrestin and releases a transcription factor. While these modifications enhance the robustness of the Tango signaling output, the added signal peptide and codon optimization bypass the native biosynthetic pathway and the C-terminal fusion intentionally alters their signaling activity. Scalable approaches to survey other aspects of GPCR biochemistry will require a more native, versatile genetic library.

To construct a complete, assay-agnostic library that can be screened in parallel, we first collected all human GPCR transcript sequences within the Ensembl database and cross checked their amino acid sequences against the Uniprot database. We then added transcript sequences for 174 experimentally-validated GPCR splice variants. To facilitate universal immunological detection, we added an N-terminal influenza hemagglutinin (HA) epitope tag to all class A and olfactory receptors. For receptors with signal peptides (i.e. classes B, C, and F), we instead inserted the HA tag between the predicted signal peptide cleavage site and the structured region of the N-terminal domain, as predicted by SignalP (see *Methods*). To facilitate the characterization of these receptors in parallel, we incorporated an attB recombination site and a 10-base unique molecular identifier (UMI) sequence upstream of the ORF (Fig. 1A); two common features used in deep mutational scanning workflows.^21–23^ The attB site facilitates the generation of stable cells that express a single receptor from a defined genomic locus while the UMI enables sequencing-based identification of receptors in the downstream assay, as is described further below. Importantly, the genomic recombination of this plasmid installs the GPCR-encoding ORF downstream of a tet-inducible promoter, which ensures that only recombined receptor genes are expressed in response to doxycycline (Fig. 1A).^21^ Together, this collection of plasmids can be used to efficiently generate stable cell lines that inducibly express a single receptor of interest or a pooled cellular library that collectively expresses a comprehensive collection of receptors.

**Figure 1.**
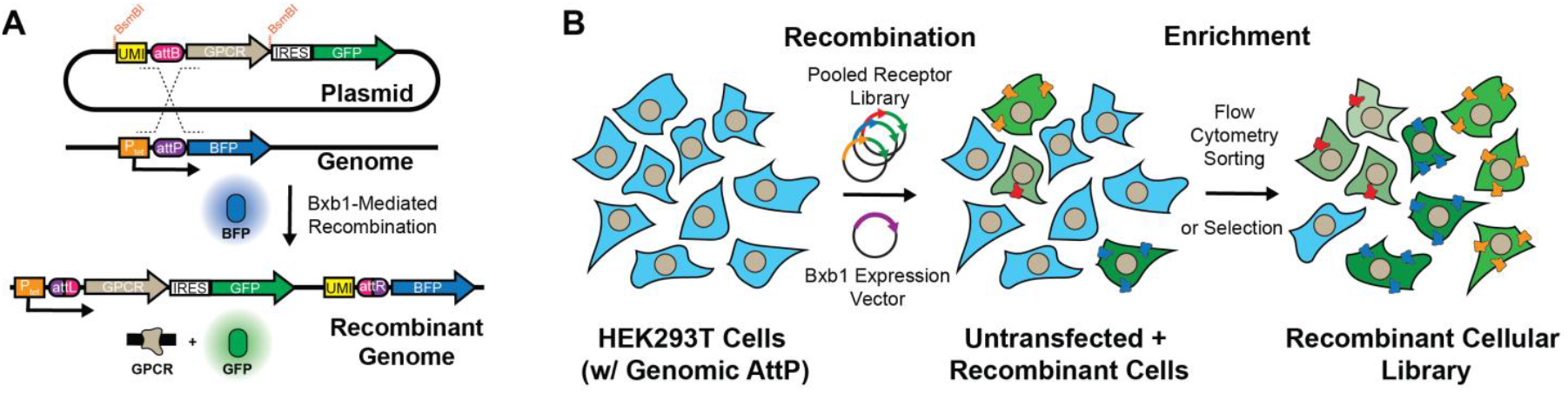
Generation of Pooled Recombinant Cellular GPCR Libraries. Cartoon schematics depict the design of the genetic elements, the manner GPCR cDNA are recombined into the genome, and how this can be used to generate a pooled cellular library in which each cell inducibly expresses a single receptor from the library. A) Plasmids carrying a single GPCR cDNA and their corresponding unique molecular identifier (UMI) sequences undergo irreversible Bxb1-mediated recombination between their attB sites and a genomic attP site. Recombinant cells stimulated with doxycycline lose BFP expression and gain bicistronic expression of the GPCR and GFP. B) Cotransfection of a genetically modified HEK293T cell line bearing a single genomic attP “landing pad” are cotransfected with a pool of GPCR-encoding plasmids and a Bxb1 recombinase expression vector. Recombination between these plasmids and the genomic landing pad in the following days generates a mixed pool of parental cells (BFP+) and recombinant cells (GFP+/ BFP-) expressing individual GPCRs from the pooled library. Recombinant cells are then enriched by FACS in order to generate the recombinant cellular library.

Pooled workflows are potentially compatible with numerous fluorescence or selection-based assays that can be used to track the expression, localization, and/ or signaling output of these receptors in parallel. To ensure this library is compatible with a variety of assays, we added BsmBI cleavage sites upstream of the UMI and at the immediate 3ʹ of the ORF to facilitate the transfer of this library from one plasmid backbone to another with a single golden gate assembly reaction (Fig. 1A). To ensure ORFs are not altered in the cloning process, we silenced native BsmBI sites where necessary and added a single 3ʹ glycine codon to each transcript (see *Methods*). Beyond these modifications and the 5ʹ epitope tag, we elected to maintain the native nucleotide sequence encoding each receptor. As a starting point, we incorporated these designs into a plasmid backbone containing a 3ʹ IRES-GFP cassette that marks recombinant cells with GFP expression (Fig. 1A). Our final library encodes 940 receptors, including 286 class A, 30 class B, 14 class C, 11 class F, and 393 olfactory, 27 taste, 4 vomeronasal receptors, 1 unknown as well as 174 splice variants (Doc. S1).

### Identification of Unstable GPCR Transcripts

To establish a one-pot approach to characterize these receptors, we generated a cellular library in which each cell expresses a single receptor by recombining this plasmid pool into an HEK293T cell line bearing a unique genomic attP site. We then used flow cytometry-based cell sorting to isolate GFP+/ BFP-recombinant cells (Figs. 1B). Deep sequencing of the UMI region within the recombined cells confirmed that 930 of the receptors encoded in the plasmid library are present in the recombined pool, with ~99% of GPCRs contributing at least 0.01% of the pooled receptor library (Fig. S1). However, in contrast to recombinant cells expressing a single receptor, the recombinant cellular library exhibited wide variation in IRES-GFP intensities (Fig. 2A). Given that cells within this pool express each receptor from an identical promoter at a single genomic locus (Fig. 1B), this variability suggests there may be differences in the cellular accumulation of the transcripts encoding these receptors. To identify receptors expressed in cells with attenuated IRES-GFP levels, we used fluorescent cell sorting to separate the low- and high-GFP cells within the library, then used deep sequencing to compare the relative abundance of the recombined receptors within each subpopulation. Our results reveal that there are 377 of 756 canonical transcripts that are statistically enriched within the low-GFP subpopulation (Fischer’s Exact Test, *p* < 0.05), some of which are enriched by up to 27-fold. Notably, 78% of these low-GFP transcripts are olfactory receptors while those encoding class A, B, C, or F receptors are primarily found in the high-GFP population (Doc. S1, Fig. 2B). To confirm that the low IRES-GFP intensity arises from low transcript levels, we measured the relative abundance of each receptor-encoding transcript by RNA-Seq (Doc. S1). When adjusting for the relative abundance of the recombined receptors in the cellular pool, we find the relative abundance of the expressed transcripts to be anticorrelated with the enrichment of recombined cDNAs within the low-GFP population (Spearman ρ= −0.697, *p* = 1.32 × 10^−111^, Fig. 2C). Together, these results strongly suggest the coding regions of certain transcripts bear features that modulate their accumulation within the cell.

**Figure 2.**
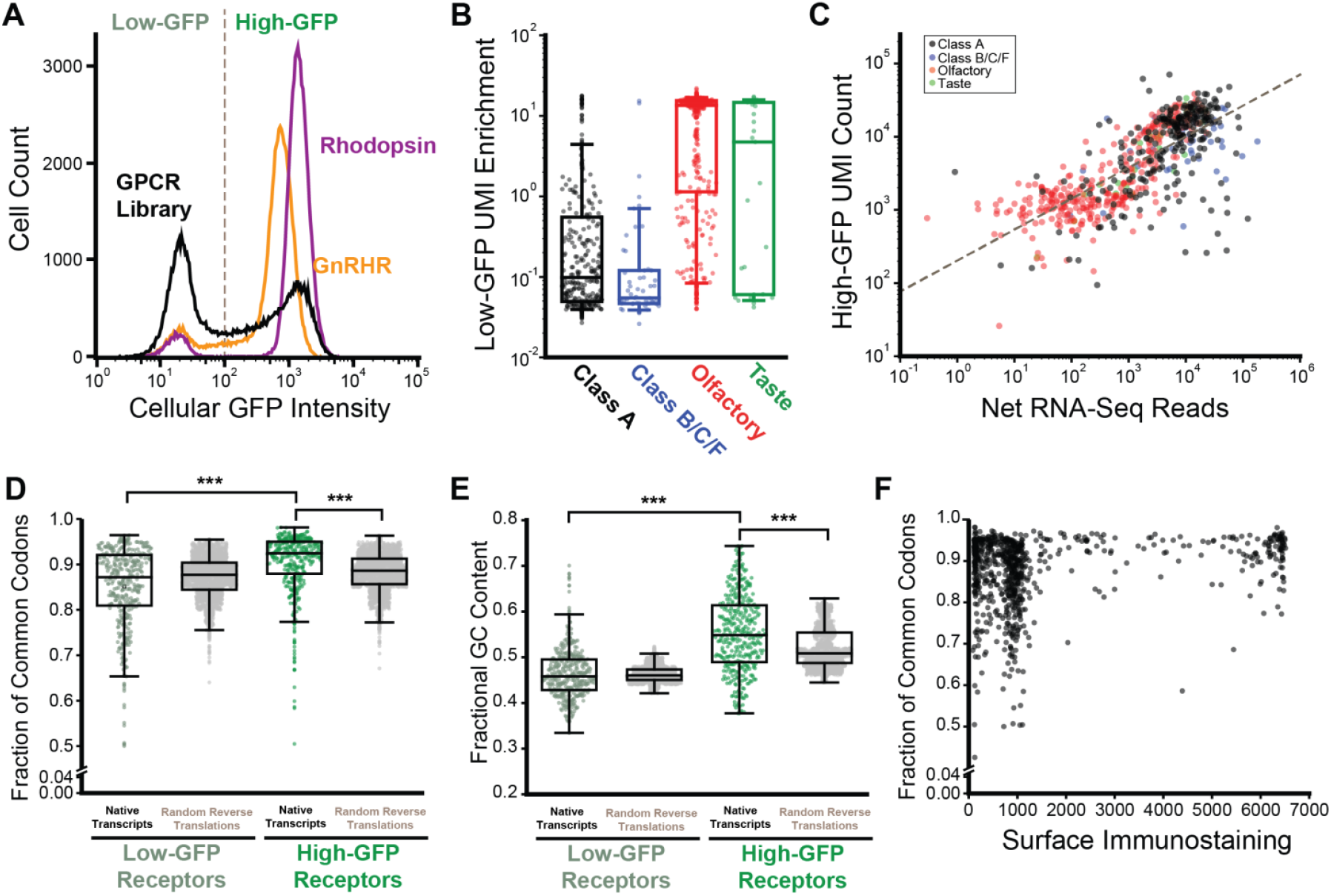
Trends in the Relative Abundance of GPCR Transcripts. The relative abundance of GPCR transcripts are characterized in the context of a pool of recombinant HEK293T cells that each express a single receptor from the GPCR library. A) A histogram depicts the distribution of cellular GFP intensities among HEK293T cells that have been recombined with the rhodopsin plasmid (purple), the GnRHR plasmid (orange), or the pooled GPCR plasmid library (black). A gray dashed line depicts the approximate position of the flow cytometry gate used to separate the high- and low-GFP subpopulations. B) A box and whisker plot depicts the degree to which recombinant cells expressing class A receptors, class B/C/F receptors, olfactory, or taste receptors are enriched within the low-GFP subpopulation. C) The extent to which recombinant cells expressing each receptor are enriched within the low-GFP subpopulation is plotted against the number of RNA-Seq reads that map within their corresponding transcript across the whole population. D) A box and whisker plot depicts how the fraction of common codons varies among transcripts that are enriched within the low- or high-GFP sub-populations. Corresponding distributions for random reverse translations of the corresponding protein sequences are shown as a reference null distribution. E) A box and whisker plot depicts how the fractional GC content varies among transcripts that are enriched within the low- or high-GFP sub-populations. Corresponding distributions for random reverse translations of each protein sequence are shown as a reference null distribution. Brackets in D) and E) reflect statistically significant differences according to a Mann-Whitney U-test (*** p < 0.0001). F) The fraction of common codons for each transcript is plotted against the corresponding plasma membrane expression of its corresponding GPCR protein.

We evaluated a variety of sequence elements to identify features that may negatively regulate transcript abundance. We first evaluated codon usage using %MinMax^24^ and found that transcripts enriched within the high-GFP subpopulation have a higher fraction of common codons relative to those within the low-GFP sub-population (Fig. 2D). Notably, common codon usage in these transcripts is also higher than would be expected relative to their genome-wide codon usage frequencies (Fig. 2D), suggesting they may have evolved to maximize translation efficiency. High-GFP transcripts also have elevated G-C content (Fig. 2E), which may help them to evade certain mRNA decay pathways.^25^ Taken together, these observations suggest high GC transcripts that rely on commonly used synonymous codons tend to accumulate at higher levels in a manner that has a clear influence on translational output (see *Discussion*).

### Identification of GPCR Transcripts with Anomalous Translation

The structural features of these transcripts that modulate their abundance may also influence their translation. To survey the translational activity of GPCR transcripts, we employed Ribo-Seq to characterize the occupancy of GPCR-encoding transcripts. Briefly, a cellular library expressing the subset of the 766 canonically spliced GPCR transcripts was treated with cycloheximide in order to arrest the ribosomes that are actively translating these receptors. Cells were then lysed and the untranslated regions of the transcripts were digested by RNAse prior to the isolation and deep sequencing of the ribosome-protected fragments. Overall, we find most GPCR-encoding transcripts have modest coverage in the context of the recombinant cellular pool. This is perhaps unsurprising, as membrane proteins are generally under-represented within Ribo-Seq data. Nevertheless, we mapped 17,178 ribosome-protected fragments to GPCR-encoding transcripts. To compare the translational dynamics across GPCR subtypes, we carried out a meta-analysis in which we grouped reads according to receptor class and binned them based on their position relative to the start and stop codons within their transcript. Each receptor class features an abundance of monomeric ribosomal footprints (monosomes, 28-38 nt) near the 5ʹ of the transcript (Fig. 3A). Downstream monosomes are evenly distributed across class A GPCR transcripts but appear to accumulate at certain positions of transcripts encoding olfactory receptors and classes B, C, and F receptors (Fig. 3A). Our experimental preparation also recovered numerous dimeric ribosomal footprints (disomes, 50-70 nt), which potentially reflect collided ribosomes. These complexes also appear to accumulate at certain points within transcripts encoding olfactory receptors and classes B, C, and F GPCRs (Fig. 3), which potentially suggests the translation of these receptors may be less processive. Notably, the occupancy of individual transcripts varies widely. Many transcripts have little to no coverage, which precludes a systematic analysis of translation efficiency across the library. Nevertheless, we do identify several transcripts that have anomalously high translational activity (Fig. 3C). Interestingly, six of the eight transcripts with the highest number of ribosome-protected fragments are olfactory receptors, four of which have below average transcript abundance (Fig. 3C). While disomes only account for 8% of the total reads, there are a few transcripts for which disomes are more abundant than monosomes such as the gamma-aminobutyric acid type B receptor subunit 1 (GABBR1). Overall, these findings reveal tremendous heterogeneity in the translational activity of GPCR-encoding transcripts.

**Figure 3.**
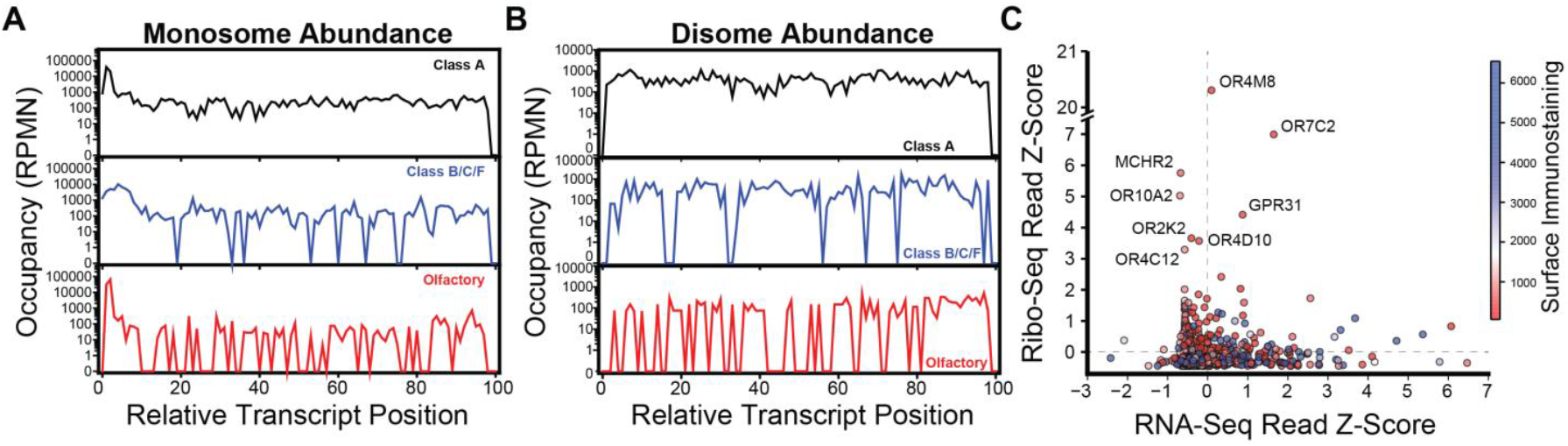
Trends in the Translational Dynamics of GPCRs. Ribo-Seq was used to compare the translational activity of GPCR transcripts in the context of a pool of recombinant HEK293T cells that each express a single receptor. Line plots depict meta-analyses of the A) monosome and B) disome pileups across subsets of transcripts encoding class A (black), or classes B/C/F (blue), or olfactory receptors (red). The number of ribosome-protected fragments per million reads per nucleotide is plotted against their relative positions within the transcript, where the codons are grouped into 100 even bins beginning with the start codon (position 0) and ending with the stop codon (position 1.0) of each transcript. C) Z-scores were calculated as a measure of how far above or below average each transcript is with respect to its coverage within the Ribo-Seq data (i.e. total ribosome protected fragments) and plotted against the Z-score for the coverage of the corresponding transcript in the RNA-Seq data (i.e. total mapped reads). Positive scores indicate the number of standard deviations above average, while negative scores indicate the number of standard deviations below average. Each point is colored according to the surface immunostaining intensity of the corresponding receptor protein. The names of outliers with high translational activity are indicated for reference.

### Plasma Membrane Expression of GPCR Subtypes

To engage in cellular signaling, most GPCRs must traverse the secretory pathway and reach the plasma membrane. To survey the plasma membrane expression of these receptors, we generated a recombinant pool of HEK293T cells expressing the full GPCR library then used a fluorescent anti-HA antibody to stain the receptors expressed at the cell surface (Fig. 4A). As expected, the sub-population of low-GFP cells that fail to accumulate receptor transcripts exhibit minimal surface immunostaining (Fig. 4B). In contrast, the surface immunostaining intensities of the high-GFP population vary by four orders of magnitude (Fig. 4B). Approximately ~64% of this high-GFP population still exhibits modest surface expression levels on par with that of cells expressing the gonadotropin hormone releasing receptor (GnRHR, Fig. 4B); a poorly expressed receptor that is known to be unstable. The remaining ~36% of this population exhibits moderate to high expression levels that are more comparable to the highly stable rhodopsin GPCR (Fig. 4B). Notably, codon usage varies widely among transcripts encoding poorly expressed receptors while common codons are abundant in transcripts encoding highly expressed GPCRs (Fig. 2F), suggesting efficient translation may be necessary to achieve the highest levels of expression. Overall, these results suggest that, while many of these receptors exhibit poor surface expression, only some of this can be attributed to variations in transcript abundance.

**Figure 4.**
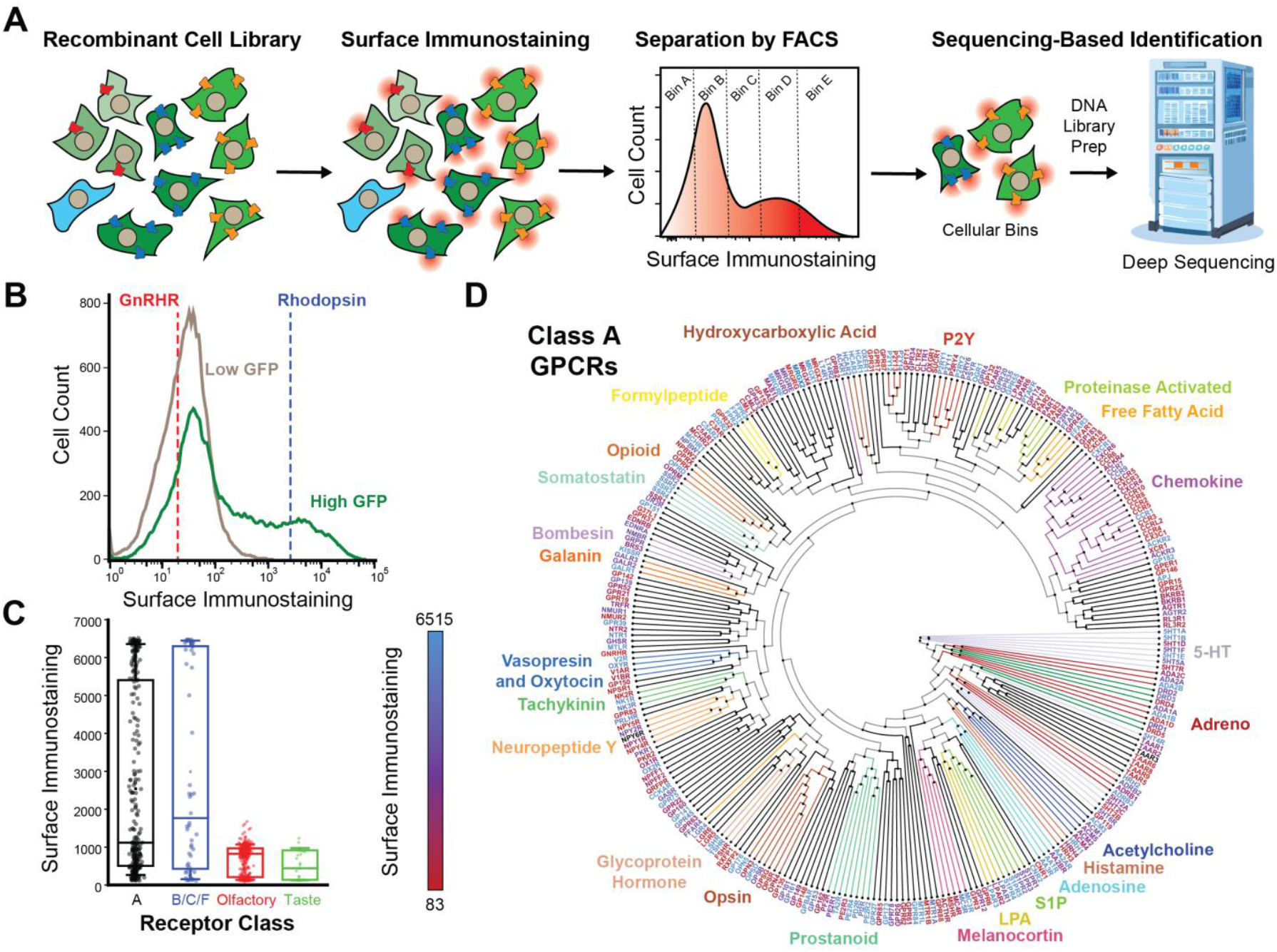
Trends in the Plasma Membrane Expression Levels of GPCRs. Deep receptor scanning was used to compare the relative surface immunostaining of 761 GPCRs in HEK293T cells. A) A schematic depicts the experimental workflow for deep receptor scanning. B) A histogram depicts the distribution of GPCR surface immunostaining intensities among the low-GFP (gray) and high-GFP (Green) subpopulations within a pool of recombinant cells that each express individual receptors from the GPCR library. The mean immunostaining intensities of recombinant cells expressing GnRHR only or rhodopsin only are shown for reference. C) A box and whisker plot depicts the distribution of surface immunostaining intensities among class A receptors in relation to those of classes B/C/F, olfactory, or taste receptors, as determined by deep receptor scanning. D) Class A receptors are arranged according to their phylogenetic relationships. The names of certain families of receptors and their corresponding branches are grouped by color. The names of each receptor are colored according to their plasma membrane expression as was determined by deep receptor scanning. Values represent the average of three biological replicates.

To further explore the diverse expression of these receptors, we adapted deep mutational scanning methodology to quantitatively compare the plasma membrane expression of each of these receptors in parallel (Fig. 4A). Briefly, we used a cell sorter to fractionate the high-GFP sub-population into five bins according to their relative surface immunostaining intensity. We then expanded each fraction and extracted their genomic DNA prior to amplification of recombined region of the genome. Finally, we employed deep sequencing to track the relative abundance of each receptor UMI across each fraction and used these data to estimate the immunostaining intensity of each receptor as previously described (Doc. S1). While class A, B, C and F receptors exhibit wide variations in expression, we note that the vast majority of taste, vomeronasal and olfactory receptors exhibit minimal surface immunostaining (Fig. 4C). A projection of immunostaining intensities onto a phylogenetic tree reveals many closely related receptors that exhibit wide variations in expression (Fig. 4D). For instance, the plasma membrane expression of the CXCR1 chemokine receptor is 22-fold higher than that of the CXCR2 receptor, which shares 77% sequence identity. Assuming they retain divergent expression profiles within neutrophils, the divergent stoichiometry of these receptors may represent a key determinant of their native oligomerization states and signaling activities.^26^ We note that receptors with the highest surface immunostaining generally have abundant transcripts, though many poorly expressed receptors are also encoded by abundant transcripts (Fig. S2). Thus, stable transcripts generally appear necessary, yet insufficient to ensure receptors accumulate at the plasma membrane. In contrast, the density of ribosomes on each transcript appears to have little connection to plasma membrane expression, as many of transcripts with the highest Ribo-Seq coverage have low plasma membrane expression (Fig. 3C). Overall, these findings suggest that while transcript features can constraints on GPCR expression, there are likely to be relatively subtle structural differences between that give rise to profound differences in their accumulation at the plasma membrane.

Our expression scan reveals tremendous heterogeneity in plasma membrane expression, which could potentially complicate the adaptation of this platform for high throughput measurements of GPCR signaling. To determine whether the poor expression of certain receptor is likely to compromise their functional characterization in the context of pooled assays, we generated five recombinant cell lines that express individual receptors in order to compare the activity of recombinant GPCRs with divergent plasma membrane expression. Using a calcium-sensitive fluorescent dye (Fura Red™), we measured the signaling response generated by a series of agonists for each corresponding receptor. Overall, we observe subtle decreases in the magnitude of the signaling responses generated by low-abundance receptors (Fig. S3). Nevertheless, we observe activation signals that are well above baseline in all cases. Based on these observations, we suspect similar pooled workflows should be feasible for fluorogenic signaling assays.

### Identification of Splicing Events that Modulate Expression

While they have yet to be structurally characterized, many of the splice variants within this library feature large insertions or deletions alter receptor structure in a manner that could change their expression and/ or function. Prior to this investigation it was unclear which of these isoforms reach the plasma membrane; a prerequisite for the activity of receptors that primarily signal from the cell surface. To determine how splicing impacts transcript levels, we evaluated whether cells expressing alternatively spliced GPCR transcripts exhibit distinct enrichment within the low-GFP subpopulation of recombinant cells expressing unstable GPCR transcripts (see Fig. 2 A-C, Doc. S1). A scatter plot suggests alternative splicing typically does not dramatically alter the enrichment patterns in cases where the canonical receptor is depleted from the low-GFP subpopulation (bottom left cluster, Fig. 5A). In contrast, splice variants of receptors that accumulate within the low-GFP population exhibit wide variations in enrichment (Fig. 5A), suggesting these modifications may tune the abundance of unstable GPCR transcripts. To determine how splicing impacts plasma membrane expression, we evaluated the surface immunostaining of each isoform in relation to its corresponding canonical receptor. Our results reveal that 133 of the 174 splice variants within this library exhibit little to no detectable surface immunostaining (Fig. 5B, Doc. S1). Nevertheless, we do identify 65 splice variants that exhibit similar expression to their canonical receptors as well as 20 isoforms that exhibit at least 20% higher surface expression relative to the canonical isoform (Doc. S1). While most such gains are modest, alternative splicing dramatically enhances the expression of four different receptors, including CCR2 (~4.5-fold), CNR1 (~6-fold), GPR132 (~7.5-fold) and ADGRE3 (~12-fold, Doc. S1). Overall, there appears to be no clear relationship between the size of the insertion or deletion and its corresponding change in transcript abundance and/ or surface expression (Fig. S4). While the functional relevance of these observations remains unclear, these results suggest some splicing modifications could potentially alter signaling outputs by modifying the stoichiometric ratios of receptors relative to certain transducers.

**Figure 5.**
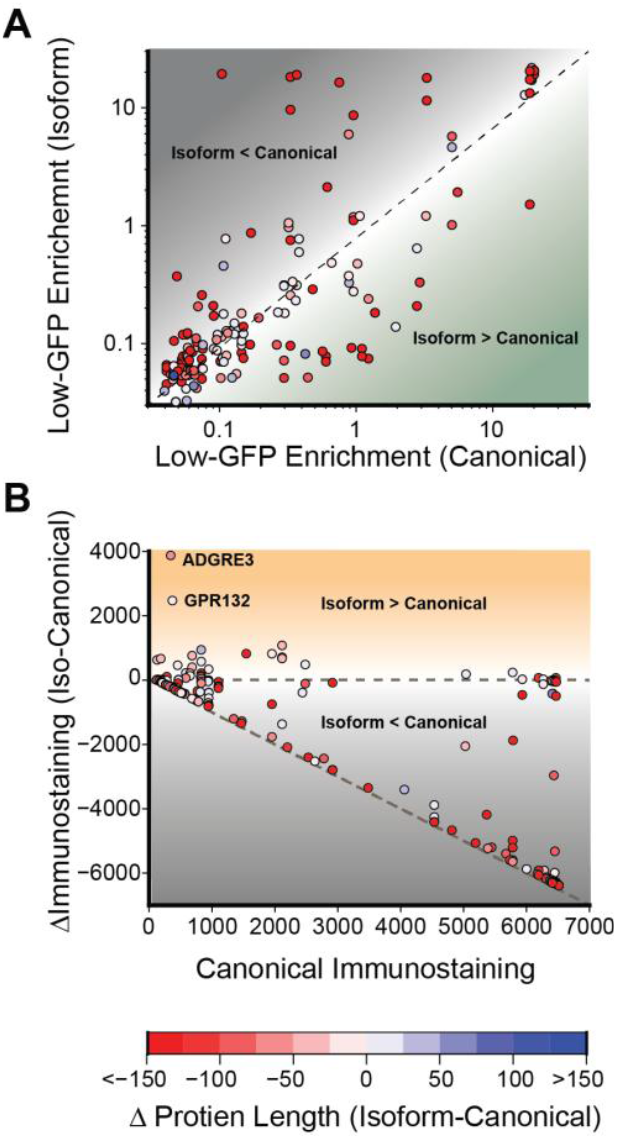
Trends in the Plasma Membrane Expression of GPCR Splice Variants. Scatter plots depict the impacts of splicing variation on receptor transcript levels and plasma membrane expression. A) The degree to which each isoform is enriched in low-GFP cells is plotted against that of the corresponding canonical receptor. Isoforms that fall above the diagonal are likely to destabilize the transcript while those that fall below are likely to stabilize the transcript based on the activity levels of the IRES-GFP cassette. B) The change in the surface immunostaining intensity of each isoform is plotted against the intensity of the corresponding canonical receptor. Isoforms that on the diagonal exhibit a complete loss of expression while those that fall in between the two dashed lines exhibit a partial loss of expression. Those above the horizontal line enhance receptor expression. Points are colored according to the change in the length of each isoform relative to the length of the corresponding canonical receptor.

### Identification of Features that Limit GPCR Expression

A previous analysis of GPCR sequence constraints suggested certain structural features may be of general importance for their expression.^27^ To identify molecular traits that modulate expression, we curated a set of features describing the characteristics of each receptor’s transcript, topology, and native structure (Doc. S2). Given the variations in GPCR transcript abundance (Fig. 2), we gathered a set of 35 descriptors for the nucleic acid sequences encoding each receptor. GPCR expression is also sensitive to the fidelity of cotranslational membrane integration;^28,29^ which hinges upon the proper recognition of hydrophobic segments by the translocon.^30,31^ To capture the relevant features for this process, we first employed DeepTMHMM^32^ to identify the general location of each receptor’s transmembrane domains (TMDs). We then used the ΔG predictor^33^ to refine the position and length of each segment as well as to estimate the translocation efficiency for each TMD (ΔG_app, pred_, see *Methods*). Finally, to capture differences in the conformational stability of the mature forms of these receptors, we extracted structural descriptors for each receptor from the AlphaFold database (canonical receptors) and a series of novel AlphaFold2 models (isoforms, see *Methods*). Together, this versatile, assay-agnostic feature set could be used to find trends in a variety of receptor scanning data sets.

To identify key determinants of GPCR expression, we employed machine learning to survey the features associated with trends in receptor immunostaining. We first employed uniform manifold approximation projection (UMAP) to survey the properties of these receptors in lower dimensional space. Our results suggest that this modest set of 75 features is generally sufficient to differentiate between the different classes of receptors (Fig. 6A). To identify features that are most closely associated with differences in plasma membrane expression, we trained a series of supervised learning models to classify receptors based on their expression levels (see *Methods*, Table S1). We first used our complete set of features to train a series of models to classify these receptors according to their plasma membrane expression of large set of 781 receptors for which each feature could be calculated (i.e. contains all seven TMDs). Our top-performing random forest model achieves considerable prediction power given the limited size of the feature set (AUROC = 0.810, Fig. 6B). However, the interpretation of feature performance is confounded by the model’s reliance on combinations of features related to properties of the mRNA transcripts and GPCR proteins.

**Figure 6.**
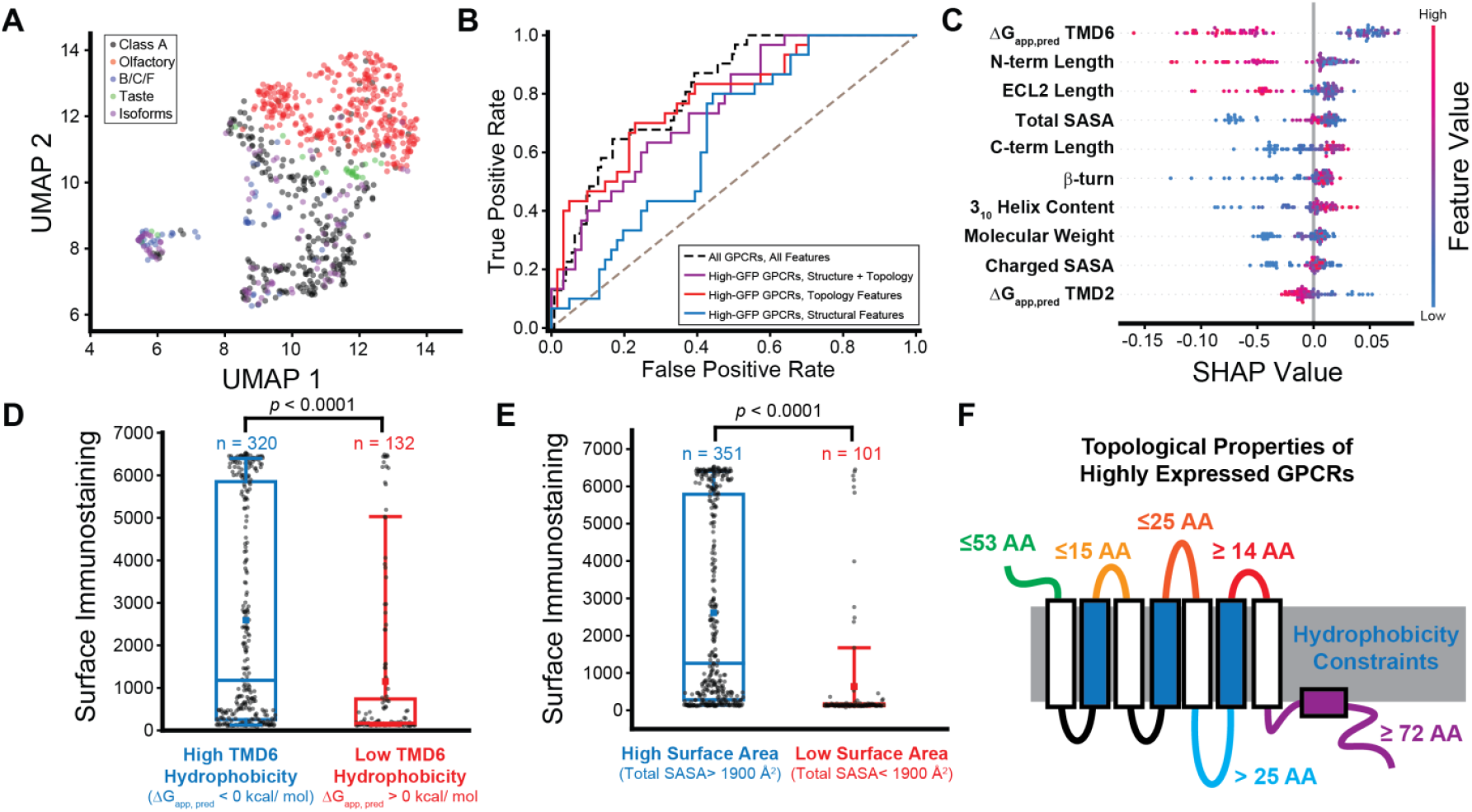
Machine Learning Analysis of Trends in GPCR Expression. Machine learning was used to identify molecular features that are associated with variations in GPCR expression. A) Uniform manifold approximation projection (UMAP) was used to reduce the dimensionality of the full feature set. The projection values for each receptor in UMAP dimension 2 are plotted against the corresponding values in UMAP dimension 1. Points are colored according to the classification of the receptors. B) A series of receiver operating characteristic curves plot the false positive rate as a function of the true positive rate for various supervised learning models including a random forest model trained on a wider set of 781 GPCRs using the full feature set (black dashes, AUROC = 0.810) as well as for random forest models trained on the expression values for the subset of 452 receptors with stable transcripts using just structural features (blue, AUROC = 0.654), just topological features (red, AUROC = 0.693), or both structural and topological features (purple, AUROC = 0.784). Models were trained to classify receptors bases on whether their surface immunostaining value is higher than the 67^th^ percentile (Intensity ≥ 2644). C) A bee swarm plot depicts the distribution of Shapley additive explanations (SHAP) values for individual receptors, which are colored according to their feature value, across the top ten most predictive features within the random forest model trained to classify high-GFP receptors based on structural and topological features. High SHAP values indicate a particular feature increases the likelihood of a low-expression classification. D) A box and whisker plot depicts the distribution of surface immunostaining intensities among receptors that feature a highly hydrophobic TMD6 (ΔG_app, pred_ <0 kcal/ mol) or a marginally hydrophobic TMD6 (ΔG_app, pred_ > 0 kcal/ mol). The two distributions are statistically distinct according to a Mann-Whitney U-test (*p* <0.0001). E) A box and whisker plot depicts the distribution of surface immunostaining intensities among receptors that have a relatively high (>1900 Å^2^) or low (<1900 Å^2^) solvent accessible surface area. The two distributions are statistically distinct according to a Mann-Whitney U-test (*p* <0.0001). F) A cartoon diagram provides an overview of the top topological properties that are associated with high expression.

To gain insights into the structural factors that limit GPCR expression, we trained an additional series of supervised learning models to classify the expression of a subset of 452 receptors that have a complete feature set and are encoded by stable mRNA transcripts (i.e. not statistically enriched within the low-GFP subpopulation, see Fig. 2A, Doc. S1). For these purposes, we used a limited set of 27 sequence-based topological features and/ or 13 features derived from models of their native structure (Table S1). Our top-performing random forest models achieve considerable predictive power regardless of whether they were trained just with structural features (AUROC = 0.654), topological features (AUROC = 0.693), or the combination of the two (AUROC = 0.784, Fig. 6B, Table S1). Interestingly, both Gini coefficients and a Shapley analysis identify the hydrophobicity of TMD6 and the total solvent accessible surface area (SASA) as the top performing topological and structural features, respectively, within the model trained using both sets of features (Table S2, Fig. 6C). Generally, receptors with marginal hydrophobicity in TMD6, which we previously found modulates evolutionary fitness,^34^ have minimal expression (Fig. 6D). Similarly, GPCRs with a total SASA of less than 1900 Å^2^ generally fail to accumulate at the plasma membrane (Fig. 6E). Overall, this survey of the top performing features reveals several topological thresholds that are associated with a loss of expression (Fig. 6F). Together, our models demonstrate that combinations of these features account for most of the observed variance in GPCR expression. While also highlighting key factors that collectively give rise to tremendous variations in the abundance of mature receptors.

## Discussion

Though advances in GPCR biochemistry have provided deep insights into the structure, function, and pharmacology of many important drug targets, many remain understudied. Indeed, approximately one quarter of non-olfactory receptors recognize unknown ligands. The development of various high throughput platforms has helped to functionally characterize some of these receptors while providing a holistic overview of GPCR specificity. However, exploring the biosynthesis, trafficking, and/ or signaling across the GPCRome will require a more versatile, comprehensive experimental platform. To this end, we have developed a genetic library encoding 940 total human GPCRs and GPCR splice variants that can be experimentally characterized in parallel using a variety of cellular assays. In conjunction with emerging cellular recombination systems, this library can be used to generate a pool of stable cells that each express a single receptor from a unique, inducible genomic promoter within a week’s time (Fig. 1). Our characterization of these pooled cell lines provides a global overview of variations in the transcript levels of individual receptors as well as systematic differences in their translational dynamics (Figs. 2–3). Using a typical deep mutational scanning workflow (i.e. Sort-Seq), we also profiled the plasma membrane expression of these receptors (Fig. 4A). Our results reveal wide variations in the expression of individual receptors and identified splice variants accumulate at the cell surface (Figs. 4–5). Finally, we curated a generalized set of descriptors for each receptor and employed machine learning to identify structural features that modulate GPCR expression. These advances provide new tools to explore GPCR biochemistry while also providing new insights into the features that influence their cellular expression.

With a single transfection, this platform can generate a population of recombinant stable cells that collectively express over 95% of known human GPCRs. These cell lines will facilitate a variety of previously unfeasible biochemical and bioanalytical experiments. Importantly, the collection of receptors encoded within the pooled library can be efficiently shuttled from one expression vector to another with a single Golden Gate Assembly reaction. This collection is therefore backward-compatible with several current activity assays and forward-compatible with future assays that couple cell sorting or cellular viability to GPCR activity, expression, or localization. Fluorogenic activity reporters can be used to survey how these receptors respond to small molecules, though the number of compounds that can be screened in this way is constrained by the limited capacity of cell sorters. Sequencing-based activity readouts could hypothetically bypass this bottleneck and facilitate high-throughput GPCRome screening against small molecule libraries, though this would require the development of economical sequencing approaches that are compatible with conventional screening equipment. Overall, cell sorting assays generally provide better signal-to-noise while sequencing based assays may offer greater scalability. Nevertheless, either signaling assay can be adapted to monitor the activation of distinct transducers by simply swapping the promoters that drive reporter expression. In addition to these “deep receptor scanning” approaches, the recombinant cells themselves may also facilitate many other biochemical and/ or bioanalytical experiments. For instance, proteomic mass spectrometry could be used to track the differential chemical modification and/ or co-immunoprecipitation of hundreds of receptors. While the exogenous expression levels and the cellular environment of these experiments is far from native and trends may be different in other cell lines, it should be noted that these experiments could potentially be carried out in other cell lines upon introduction of single genomic attP site. Regardless of this caveat, we suspect this experimental platform will open the door to a variety of creative new workflows.

While the transcription levels of the recombined ORFs should be relatively uniform (Fig. 1A), recombinant cells still exhibit considerable variation in both transcript and protein levels (Figs. 2 & 4). As we opted to preserve the native codon usage, we suspect divergent transcript levels arise from the intrinsic sequence and structural features of these transcripts. Indeed, we find clear evidence that high G-C transcripts and transcripts bearing a higher proportion of common codons accumulate at higher levels in the cell (Fig. 2). The mRNA features also give rise to variations in translational efficiency, though the occupancy of the transcript does not appear to strictly dictate protein levels (Fig. 3C). These features also alter where ribosomes accumulate within transcripts (Fig. 3 A-B), which could influence the cotranslational assembly of nascent receptors. Our data also reveal the prominent effects of splicing on the abundance of transcripts and receptor proteins. Overall, we find that 52.3% of the alternatively spliced transcripts exhibit a decrease in abundance of at least 50% relative to the canonical transcript (Doc. S1). While most splice variants exhibit little to no plasma membrane expression, including those with low expression of the corresponding canonical receptor, we identify 83 isoforms that are within 50% of the canonical surface immunostaining intensity (Doc. S1, Fig. 5). We also identify several isoforms that exhibit dramatic increases in plasma membrane expression. In the case of these of GPR132 and ADGRE3, we note that the canonical forms of these receptors exhibit little to no surface immunostaining whereas the isoforms achieve a more typical expression level (Fig. 5). Our wider analysis of expression trends also identified a variety of structural features of the receptor proteins that modulate their plasma membrane expression (Doc. S2, Fig. 6). Interestingly, our machine learning models suggest the most important topological feature is hydrophobicity of TMD6, which also undergoes considerable translocation during GPCR activation. Indeed, we recently found that evolutionary modifications to the hydrophobicity of this segment influence the cotranslational assembly efficiency of the gonadotropin releasing hormone receptor (GnRHR) in a manner that tunes its maturation. Importantly, the divergent TMD6 hydrophobicity among GnRHRs tracks with mammalian reproductive phenotypes and modulates certain epistatic interactions that shape protein evolution.^34,35^ Additional investigations are needed to explore how evolution balances these factors against those that are directly involved in signal transduction.

Beyond these discoveries, we believe this experimental platform could usher in a new era of accessible high-throughput GPCR biology. The development of our genetic library, which is freely available on Addgene (ID numbers 243681 and 243682), was facilitated by the decreasing costs of gene synthesis. Moreover, the number of DRS applications will continue to grow with the continued development of massively parallel genetic technologies. Importantly, these experiments circumvent the need for boutique robotics and only require a cell sorter, a thermocycler, and modern DNA sequencing. Advances in structural modeling will also continue to empower generalizable machine learning approaches to extract molecular insights from these experimental outputs. The models and feature set employed herein, which can be further expanded to include other descriptors, can be used to analyze any future DRS data regardless of the nature of the assay. It is important to note that the number of experimental descriptors for these receptors will also continue to increase with each additional deep receptor scan. Thus, these machine learning analyses will only become more powerful over time as training data continue to accumulate. With sufficient data, deep learning models may eventually be able to elucidate how hidden variables associated with expression levels, modification states, cellular trafficking, and transducer coupling can interact to create complex signaling behaviors. These hybrid data science approaches hold great promise for the future of receptor biology.

## Materials and Methods

### GPCR Library Design and Production

A collection of human GPCR genes were first collected based on their annotations in the Uniprot database. The corresponding canonical transcript sequences for these genes along with a list of experimentally validated isoforms^36^ were then collected from the Ensembl database. All receptors encoded by transcripts under 3 kb were included in the downstream production of the library. For GPCRs lacking an extended N-terminal domain (i.e. class A and olfactory receptors), we inserted an influenza hemagglutinin (HA)-coding sequence after the first 5ʹ start codon and the second codon in order to facilitate universal immunological detection. For the remaining receptors (i.e. classes B, C, and F), we first used TOPCONS^37^ to identify transcripts containing signal peptides then predicted their signal peptide cleavage site using SignalP 6.0^38^ and the positions of the structured residues within the downstream N-terminal domain using NetSurfP-2.0.^39^ We then designed a series of HA-tagged versions of each signal peptide-containing receptor by inserting the tag residues at different positions between the predicted cleavage site and the beginning of the structured N-terminal residues. The signal peptide cleavage site and the ordering of the N-terminal residues was then predicted again for each design, and the construct that was most strongly predicted to preserve the correct cleavage site without altering disorder prediction was chosen as the final design for each receptor.

To facilitate robust sequence-based identification, we developed a custom algorithm that generated a set of maximally differentiable 10-base UMI sequences. A custom python script was first used to generate all possible 10N nucleotide sequences that have a minimum Hamming distance of 4, a maximum homopolymer length of 2, and were restricted to having a GC content between 35% and 65%. UMI sequences containing common restriction sites and/ or ATG codons were then removed from the set. The remaining UMI sequences were then randomly assigned to individual GPCR transcripts. We then appended a cassette containing a BsmBI cleavage site, a single UMI, an attB recombination site from 5ʹ to 3ʹ, respectively, upstream of the start codon of each coding sequence. To facilitate Golden Gate assembly, we also replaced the native stop codon of the coding sequence with a glycine codon, a stop codon, and an additional BsmBI cleavage site. All native C-terminal glycine codons were removed in order to prevent the introduction of 44 non-native diglycine motifs. Finally, to prevent any extraneous cleavage events during cloning, we introduced silent mutations to the most common codon possible at 235 native BsmBI sites within coding regions. Each coding set was then incorporated upstream of an IRES-GFP cassette in the context of a modified pcDNA5 backbone that has been used for previous deep mutational scanning experiments.^40^ A complete map of each plasmid can be found in the accompanying Mendeley Data directory (doi:10.17632/3b4n36z4bg.1).

A final collection of 940 individual plasmids, which includes 766 encoding canonical receptors and 174 encoding isoforms, were synthesized and sequenced by the GenScript Biotech Corporation (Piscataway, NJ). Plasmids encoding each of the 940individual receptors were suspended in water and stoichiometrically pooled prior to electroporation of the plasmid library into electro-competent NEB10β cells (New England Biolabs, Ipswitch, MA). Electroporated cells were then grown in liquid culture overnight and purified using the ZymoPure endotoxin-free midiprep kit (Zymo Research, Irvine, CA).

### Cell Culture

These investigations were carried out using a previously described HEK293T cell line^21^ that contains a single genomic “landing pad” featuring a Tet-ON promoter, an attP recombination site, and a TagBFP gene, which were a generous gift from Douglas Fowler (University of Washington). Cells were grown in Dulbecco’s modified Eagle’s medium (Gibco, Grand Island, NY) supplemented with 10% fetal bovine serum (Corning, Corning, NY) and a penicillin (100U/ml) and streptomycin (100µg/ml) antibiotic cocktail (Gibco, Grand Island, NY) in a humidified incubator containing 5% CO2 at 37°C. Recombination was initiated by cotransfecting cells with a Bxb1 expression vector (pCAG-NLS-HA Bxb1) in combination with the GPCR plasmid pool using Fugene 6 (Promega, Madison, WI). The cells were then incubated at 33° C for 3 days post-transfection. GPCR expression was induced by the addition of 2 µg/mL doxycycline to the media beginning two days after transfection. The cells were then incubated at 37° C for 24 hours prior to the isolation of 2M+ recombinant BFP-negative cells using a Bigfoot Spectral Cell Sorter (Thermo Fisher Scientific, Waltham, MA USA). The resulting enriched pool of recombinant cells were grown in 10 cm dishes with complete media supplemented with 2 µg/mL doxycycline until they reached confluency, at which point they were divided into 15 cm dishes for the completion of experimental replicates.

### Fluorescence Activated Cell Sorting

Surface immunostaining and GFP intensities of recombinant cells were analyzed 7-12 days post-transfection. Briefly, the adherent cells were washed with phosphate buffered saline (AthenaES, Baltimore, MD) and harvested with 0.25% Trypsin-EDTA (Gibco, Grand Island, NY). HA-tagged GPCRs expressed at the plasma membrane were immunostained by incubating the cells with a DyLight550–conjugated anti-HA antibody (Thermo Fisher, Waltham, MA) for 30 minutes. Cells were then washed twice with phosphate buffered saline and passed through a 40 μm cell strainer to remove cellular aggregates prior to separation with a Bigfoot Spectral Cell Sorter (Thermo Fisher Scientific, Waltham, MA). A series of hierarchical gates were set to first divide cells into two populations based on their GFP intensity (see Fig. 2A). The high-GFP subpopulation was then separated into quintiles based on Dylight550 intensity further fractionated into quintiles. At least 800,000 cells from the low-GFP subpopulation and 600,000 cells from each high-GFP quintile were isolated to ensure deep sampling. Fractionated sub-populations were each expanded in 10 cm culture dishes prior to harvesting and freezing 15-25 million cells per subpopulation for the downstream genetic analysis.

### Extraction and Sequencing of Cellular DNA and RNA

To quantify the relative abundance of recombined cells expressing each receptor within each cellular isolate, we first extracted genomic DNA from each cellular sub-population using the DNeasy Blood & Tissue Kit (Qiagen, Hilden, Germany). A previously described semi-nested PCR technique was then used to selectively amplify the barcoded region of the recombined plasmids within the gDNA.^40,41^ Briefly, an initial PCR reaction was used to amplify the recombinant region of the genomic DNA. This PCR product was then used as a template for a second PCR reaction that amplifies the barcoded region and installs indexed Illumina adapter sequences. Libraries were sequenced using an AVITI System Sequencing Instrument (Element Biosciences, San Diego, CA) for 2×75 cycles at an average depth of ~4 million reads per subpopulation.

RNA-Seq analysis was carried out on an enriched population of BFP-recombinant cells grown in complete media supplemented with 2 µg/mL doxycycline. Cells were grown to confluency in 10 cm dishes (10-20 million cells) prior to the extraction of total cellular RNA using an Aurum™ Total RNA Mini Kit (Bio-Rad Laboratories, Hercules, CA). RNA was further purified using RNAClean XP Bead Based Reagent (Beckman Coulter, Brea, CA). Poly A+ libraries were constructed using a Poly(A) mRNA Magnetic Isolation Molecule (New England Biolabs, Ipswich, MA) in conjunction with an xGen RNA Library Prep Kit (Integrated DNA Technologies, Coralville, Iowa) as directed by manufacturers’ instructions. Libraries were run using a 300 cycle NovaSeq X (Illumina Inc., San Diego, CA) to a final depth of at least 300M reads per sample. Final mock-normalized read counts represent the difference between the average read count among recombinant cells expressing the GPCR library and the parental cell line across three biological replicates (see Doc. S1).

### Deep Receptor Scanning Calculations

Deep sequencing data were analyzed in a manner similar to those described previously described.^40,41^Briefly, we first removed reads that were likely to contain more than one error based on Q-scores as well as any reads that do not contain an exact match for one of the 940GPCR UMIs. The remaining reads from each cellular subpopulation were then rarified down to a common number of reads equal to the sample with the lowest total reads. The degree to which recombinant cells expressing each individual receptor are enriched within the low-GFP was then determined by dividing the total number of reads for each receptor in the low-GFP population by the number of corresponding reads across the five high-GFP subpopulations combined. Enrichment scores were only calculated for receptors that were sampled at a frequency of at least 0.0005% within each subpopulation (i.e. 10 reads for low GFP, 50 reads for high GFP). Surface immunostaining levels were then estimated by calculating the immunostaining intensity-weighted average of the reads for each receptor across the five high-GFP subpopulations. Receptor intensities from each replicate were normalized relative to one another using the overall mean surface immunostaining intensity of the recombinant cell population on each day in order to account for any small deviations in calibration. Intensity values reported herein represent the average normalized intensity values from three replicates.

### Codon Usage and Nucleotide Analysis

Synonymous codon usage in HA-tagged GPCR transcript sequences were evaluated using the %MinMax algorithm.^24,42^ Briefly, %MinMax uses a sliding window function to analyze relative codon usage over the length of an input transcript sequence. To analyze each coding sequence, the fraction of the sequence using rare or common codons was calculated as the fraction of sequence with %MinMax <0 or %MinMax >0, respectively. Codon usage data for *H. sapiens* was sourced from the HIVE-CUT database.^43,44^ The %MinMax output in Figure 2 was calculated using a sliding window size of 17 codons, though similar trends were also observed using sliding window sizes of 7 and 27 codons. GC content was calculated as the fraction of the sequence using G or C nucleotides. The GPCRs were split into two groups based on whether the fraction of their sequencing reads in the high-GFP sub-population fell above (high-GFP GPCRs) or below (low-GFP GPCRs) the median value. The fraction of rare and common codons and GC content were then compared across the two groups.

As a null model, 200 random reverse translations (RRTs) were generated using *H. sapiens* codon usage frequencies for the top- and bottom-ten GPCRs with respect to the fraction of sequencing reads in the high-GFP sub-population.^24^ The fractions of rare and common codons in each RRT, as well as the GC content, were then calculated as described above for the RRTs.

### Ribosome Profiling Analysis

The translational dynamics of recombinant cells expressing a collection of 766 canonical GPCRs was characterized using the RiboLace platform.^45^ Briefly, an enriched population of 5-10 million live recombinant cells (GFP+/ BFP-) were incubated in PBS containing 10 μg/mL cycloheximide (CHX) for 5 minutes at 37°C in order to arrest actively translating ribosomes. Cells were then harvested with a cell scraper and washed twice in cold PBS containing 20 μg/mL CHX, pelleted by centrifugation, and flash frozen at −80°C. The downstream molecular biology was carried out by Immagina Biotechnology (Povo, Italy), who extracted cellular ribosomes then isolated and sequenced the ribosome-protected fragments in accordance with their previously described protocol.^45^ Sequencing data were trimmed with cutadapt (Martin, 2011) and UMI-tools to remove Illumina linker regions and UMI sequences from each read, respectively.^46^ Non-coding RNA sequences were obtained from RNAcentral (https://rnacentral.org/) and removed from sequencing data using Bowtie2.^47^ The remaining reads were then mapped to GPCR transcript sequences using STAR.^49^ Ribosome-protected fragments ranging in size from 28-38 nt were designated as monosomes while 50-70 nt reads were designated as disomes. Reads that fell outside these ranges were discarded from the analysis. A meta-analysis of the ribosome pileups across different receptor sub-types was carried out using a custom software package (https://github.com/schlebachlab/DRS). Isoform-encoding vectors were excluded from the genetic libraries employed for these experiments due to the fact that ribosome-protected fragments within their transcripts cannot be differentiated from those within canonical transcripts.

### Signaling Assays

A calcium-sensitive fluorogenic dye was used to characterize the signaling of various receptors in recombinant HEK293T cells. Briefly, recombinant cell lines expressing individual receptors were generated as described above then isolated using fluorescence activated cell sorting 5 days post-transfection. Cells were analyzed 7-12 days after isolation. Signaling assays were carried out in 96 well plates by plating 100k cells in 100uL of media per well 24 hours prior to the experiment. Each condition was plated in triplicate for each biological replicate. Assays were carried out using the Fura 2 QBT Calcium Kit in Hank’s Balanced Salt Solution (HBSS, Molecular Devices, San Jose, CA, USA). A Fura dye solution was prepared by adding 10 mL of HBSS to a vial of component A from the Fura 2 Calcium Kit. Concentrated agonist stocks were prepared at 1.1 uM or 1.1 mM in HBSS, which were diluted to final concentrations of 100 nM and 100 uM, respectively. Prior to the initiation of measurements, 100 uL of the Fura dye solution was added to each well and incubated in the dark at 37°C for 1 hour. Two baseline readings (340 nm/ 510 nm excitation/ emission, 380 nm/ 510 nm excitation/ emission) were then collected for each well using a Synergy H4 Microplate Reader (BioTek, Winooski, VT, USA). The agonists were then mixed into each well prior to repeating the fluorescence measurements. Emission ratios were then calculated over time and corrected by subtracting the average ratio of the vehicle-treated mock sample.

### Machine Learning Features

Sequence- and structure-based descriptors for each receptor were derived from their nucleotide sequences, amino acid sequences, and from models of their native structures. Sequences from the Ensembl database were used to generate nucleotide-based descriptors of each transcript based on a set of descriptors that were previously used to predict the protein expression levels of integral membrane proteins.^48^ Protein sequences were processed using the Biopython ProtParam module to calculate physicochemical properties including molecular weight, isoelectric point, aromaticity (representing the fraction of aromatic amino acid residues), the instability index based on dipeptide composition, and the Grand Average of Hydropathy (GRAVY) score using the Kyte-Doolittle hydrophobicity scale. Protein sequences were also used to derive topological descriptors of the nascent TMDs that form during biosynthesis, which are often distinct from those that are present in the natively folded protein.^50^To gather topological descriptors for the nascent receptors, the approximate positions of the TMDs within each protein were first predicted using DeepTMHMM,^32^ a deep neural network trained to identify the topological features of membrane proteins. The portions of these TMDs that are most likely to undergo translocon-mediated membrane integration was then identified using a custom python script that scans each TMD region identified by DeepTMHMM to identify the precise segment that minimizes the transfer free energy (ΔG_app,pred_) according to the White & von Heijne biological hydrophobicity scale.^33^ The positions and lengths of TMDs and loops within each receptor were derived from these predictions. Structural descriptors of canonical receptors were derived from structural models contained within the AlphaFold Protein Structure Database (https://alphafold.ebi.ac.uk/).^51^ Structural models of each isoform we generated using the AlphaFold2^52^ module within ColabFold^53^ prior to feature extraction.

Certain topological features were normalized in a manner that improved model performance. For instance, the lengths of all relevant topological features such as TMDs, extracellular/ intracellular loops, and the N- and C-terminal tails were normalized by the length of the processed full-length protein. In addition to topological descriptors, amino acid sequences were used to derive various other general properties such as molecular weight, isoelectric point, the fraction of aromatic residues (aromaticity), a dipeptide-based instability index,^54^ and the average hydrophobicity according to the Kyte-Doolittle scale (GRAVY Score).^55^ Structural models were analyzed using FreeSASA^56^(https://freesasa.github.io/) to generate comparative descriptors of the solvent accessible surfaces of native GPCR. The total accessible charged (sasa_crg), polar (sasa_plr), and/ or apolar (sasa_aplr) surface areas were calculated for each receptor then normalized by the corresponding total surface area of the receptor to determine the fractional surface area of each class of chemical group (i.e. f_crg, f_plr, f_aplr). Finally, the fractional content of common secondary structure elements, including α-helices, 3_10_-helices, extended strands, isolated β-bridges, turns, and coils were calculated using STRIDE.^57^ A complete list of features describing these transcripts and the receptors they encode is included in Doc. S2.

### Model Training

Features were analyzed using both supervised and unsupervised machine learning approaches in order to explore trends and predict GPCR plasma membrane expression (PME). To search for clusters of receptors with similar features, we first employed Uniform Manifold Approximation and Projection (UMAP)^58^to reduce the dimensionality of the feature set. Input features were standardized using StandardScaler from scikit-learn, and UMAP embeddings were generated using default parameters (n_neighbors = 15, min_dist = 0.1, metric = ‘euclidean’). Supervised classification models were trained to differentiate between high- and low-PME receptors based on deep receptor scanning measurements. Receptors were designated as either high- or low-PME based on a cutoff at the 67th percentile (PME = 2644). Receptors were then randomly divided into 80:20 training and test set. Each training set consisted of approximately 67% low PME receptors to avoid skewing the training and/ or test distributions. We trained a series of Random Forest classifier models using four biologically motivated feature subsets including (1) the full set of features including protein and transcript features, (2) a subset of protein features including topological and structural features (3) topological features derived from sequence-based membrane topology predictions, and (4) structural features derived from the collection of AlphaFold2 models. Model performance was evaluated based on its ability to correctly classify each test set using standard classification metrics, including balanced accuracy, precision, recall, F1 score, and the area under the receiver operating characteristic curve (AUROC). SHapley Additive exPlanation values (SHAP),^59^ a game theory approach to quantify the contribution of each feature to individual predictions, were used to interpret model behavior.

## Resource Availability

### Lead Contact

All inquiries can be directed to Jonathan Schlebach (jschleba (at) purdue.edu).

### Materials Availability

The pooled GPCR expression vector libraries described herein are available through Addgene (ID numbers 243681 and 243682). Individual GPCR expression plasmids will be freely shared with the research community upon request. HEK293T cells that were used to generate recombinant cell lines were a kind gift from Douglas Fowler.

### Data and Code Availability

Code for the analysis of sequencing data associated with deep receptor scanning experiments can be found on the Schlebach Lab GitHub page (https://github.com/schlebachlab/DRS). Sequencing data for deep receptor scanning can be accessed through NCBI (PRJNA1280020). Sequencing data for RNA-Seq experiments can be accessed through the Gene Expression Omnibus (GSE304607). Sequencing data for Ribo-Seq experiments can be accessed through the Gene Expression Omnibus (GSE307296). Surface immunostaining intensity values, processed RNA-Seq read counts, Ribo-Seq footprint counts, and fold-enrichment values for each receptor can be found in Doc. S1. Structural models and all other raw experimental data have been deposited in a freely accessible Mendeley data directory (doi:10.17632/3b4n36z4bg.1).

## Acknowledgements

We thank Rebecca Voorhees for helping to provide the inspiration for these experiments and Bil Clemons for helpful input. We thank Phillip SanMiguel and the Genomics and Genome Editing Center at the Bindley Bioscience Institute for technical assistance with next generation sequencing efforts. This work was supported by funding from the National Institute of General Medical Sciences (R35152086 to J.P.S. and DP1146256 to P.L.C.).

## Author Contributions

A.T., M.G., S.S., M.H., L.M.C., K.N., P.L.C., M.M.B., A.M., C.P.K, W.C.-M., and J.P.S. contributed to the research design. A.T., M.G., S.S., M.H., L.M.C., A.B., I.A., J.M.G., B.N.C., C.P.K., and J.P.S. performed research. J.P.R., C.B.P., P.L.C., M.M.B., A.M., W.C.-M., and J.P.S contributed analytical tools and/ or other resources. A.T., M.G., L.M.C., E.F.M., P.L.C., C.P.K., W. C.-M., and J.P.S. analyzed the data. A.T. and J.P.S. wrote the paper with input from their coauthors.

