## Supplemental Materials for "Deep Receptor Scanning Reveals General Sequence Constraints on GPCR Biosynthesis"

*Corresponding Authors: jschleba (at) purdue.edu, willow.coyote-maestas (at) ucsf.edu, and cpkuntz (at) purdue.edu

**Contents:**

-Figure S1
-Figure S2
-Figure S3
-Figure S4
-Table S1
-Table S2
-Table S3

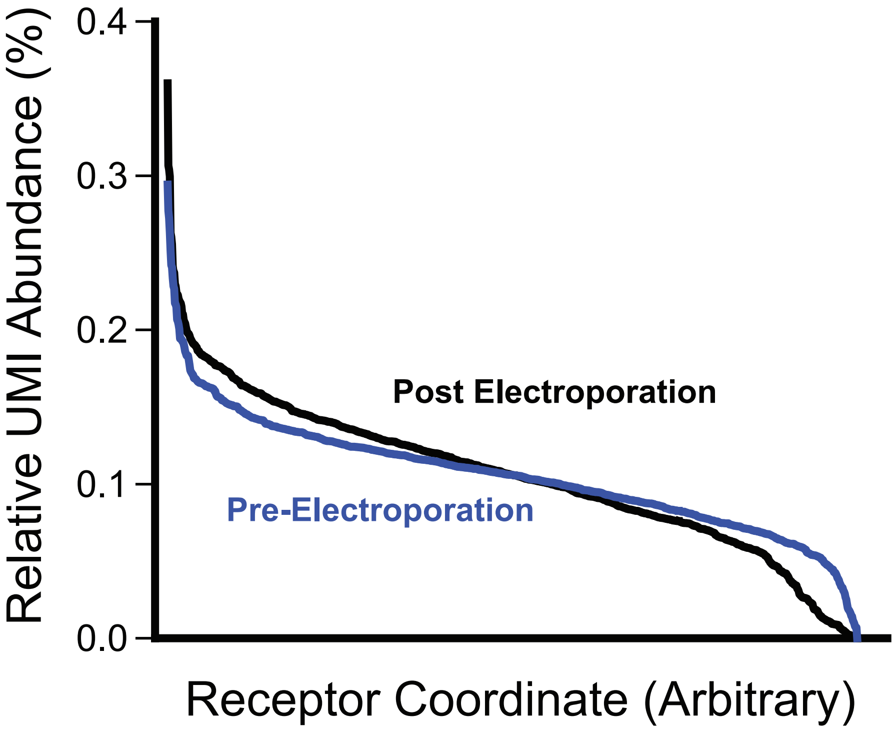

**Figure S1. Relative Abundance of GPCR Plasmids within Pooled Libraries.** 946 individual plasmid preparations were hand-mixed at an equal stoichiometric ratio prior to electroporation of the pool into *E. coli*. The plasmid pool was then purified from *E. coli* prior to comparison of the relative abundance of each GPCR-associated UMI in the initial pool and the final plasmid preparation by deep sequencing. Individual UMIs are plotted onto an arbitrary coordinate according to the percentage of the total reads withn each plasmid population. Overall, the variance between individual UMIs is typically modest and is remains similar after transformation into *E. coli*.

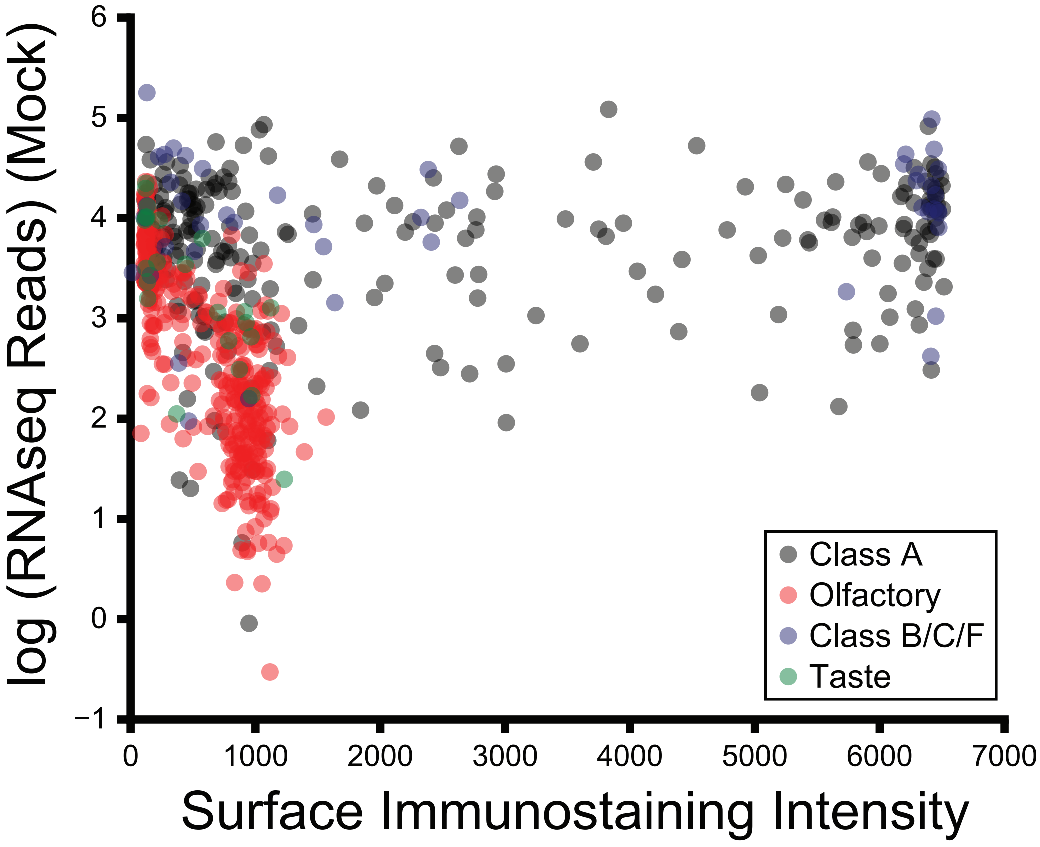

**Figure S2. GPCR Transcript Abundance and Surface Expression.** A scatterplot depicts the $\log_{10}$ transformation of the number reads per kilobase for each GPCR-encoding transcript relative to the plasma membrane expression of the corresponding receptor protein. The number pf transcript reads for each receptor within a mock transfection sample were subtracted from the number of reads for the corresponding transcript within a pool of recombinant cells expressing GPCRs in order to correct for any endogenous receptor transcripts, which should not influence the immunostaining of the HA-tagged receptors. Transcripts are colored according to the class of the receptor they encode, for reference

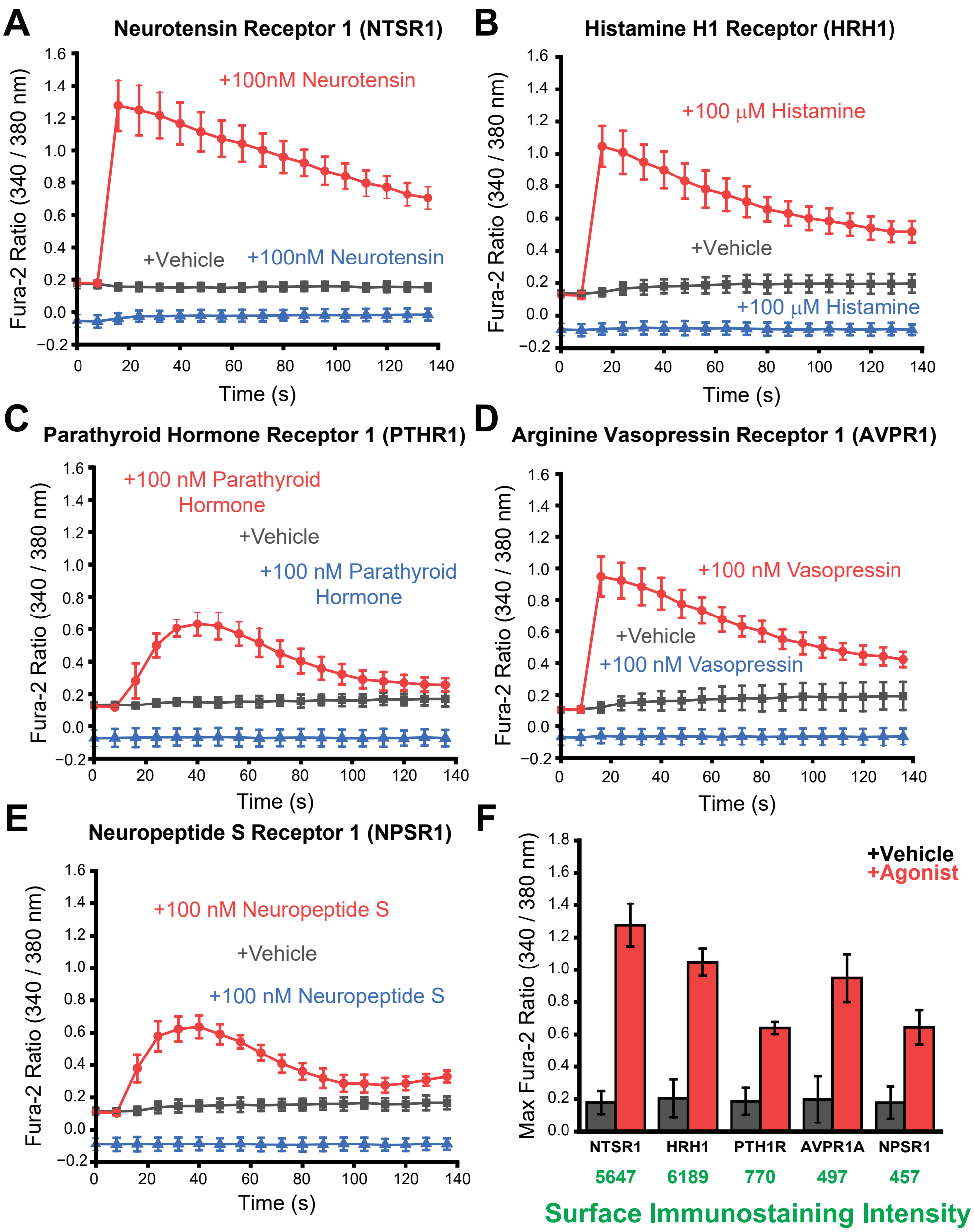

**Figure S3. Signaling Activity of GPCRs with Divergent Plasma Membrane Expression.** The signaling activity of five GPCRs with varying plasma membrane expression was compared in the context of recombinent HEK293T cells by monitoring calcium fluxes generated in the presence of agonists. A series of recombinant HEK293T cell lines expressing A) NTSR1, B) HRH1, C) PTHR1, D) AVPR1, or E) NPSR1 were incubated with a calcium-sensitive Fura-2 dye prior to introduction of agonists. A series of line plots depict the change in the Fura-2 fluorescence emission ratio over time after the addition of agonist (red) or vehicle (gray) to GPCR-expressing cells or addition of agonist to mock transfected cells (blue). F) A bar plot depicts the peak emission ratio achieved for cells expressing each receptor under each condition. The corresponding surface immunostaining intensities of each receptor by deep receptor scanning is shown in green for reference. Emission ratio values reflect the average of nine technical replicates collected across three biological replicates and error bars reflect the standard deviation. We note that the transcripts of these receptors were not observed within RNA-Seq measurements of the parental cell line. Together, these results demonstrate the the plasma membrane expression of recombinantly expressed GPCRs does not limit signal transduction in the context of this cell line.

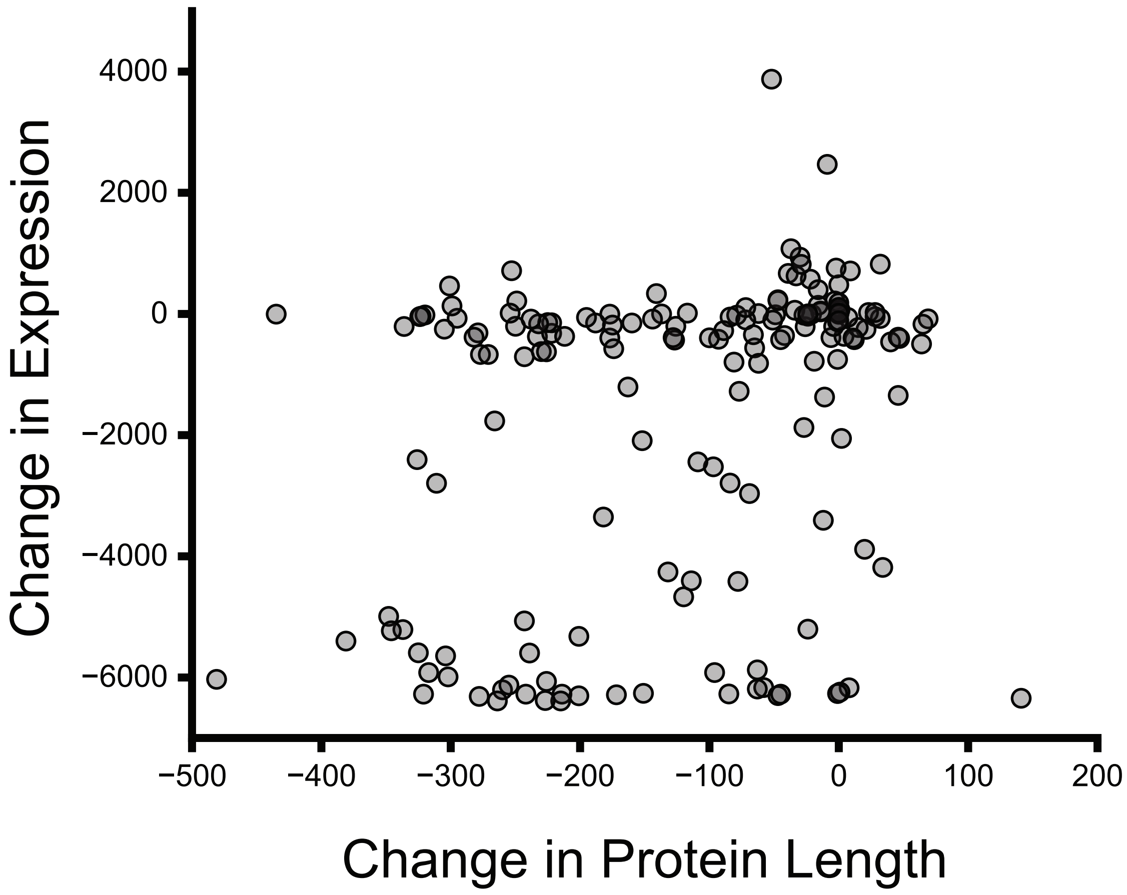

**Figure S4. Trends in the Plasma Membrane Expression of GPCR isoforms with Varying Length.** A Scatterplot depicts the change in the plasma membrane expression of each isoform in relation to its corresponding change in protein length according to deep receptor scanning measurments. Changes in expression and length are cast in terms of the isoform receptor value minus the cannonical receptor value.

### Table S1. Supervised Model Performance

*Models trained to classify the full set of GPCRs (n= 781)* w/ the full feature set (n= 72).*

| Model | Balanced Accuracy | F1-Score | Precision | Recall | ROC-AUC |
|---|---|---|---|---|---|
| Random Forest | 0.731 | 0.532 | 0.438 | 0.677 | 0.810 |
| XG Boost | 0.653 | 0.454 | 0.590 | 0.370 | 0.841 |
| Gradient Boosting | 0.596 | 0.336 | 0.455 | 0.272 | 0.828 |

*Models trained to classify GPCRs with stable transcripts (n= 452)* w/ topological and structural features (n= 40).*

| Model | Balanced Accuracy | F1-Score | Precision | Recall | ROC-AUC |
|---|---|---|---|---|---|
| Random Forest | 0.735 | 0.646 | 0.600 | 0.700 | 0.784 |
| XG Boost | 0.645 | 0.510 | 0.564 | 0.471 | 0.731 |
| Gradient Boosting | 0.617 | 0.457 | 0.538 | 0.403 | 0.710 |

*Models trained to classify GPCRs with stable transcripts (n= 452)* w/ topological features (n= 27).*

| Model | Balanced Accuracy | F1-Score | Precision | Recall | ROC-AUC |
|---|---|---|---|---|---|
| Random Forest | 0.600 | 0.415 | 0.559 | 0.337 | 0.693 |
| XG Boost | 0.624 | 0.468 | 0.555 | 0.422 | 0.704 |
| Gradient Boosting | 0.603 | 0.445 | 0.524 | 0.413 | 0.696 |

*Models trained to classify GPCRs with stable transcripts (n= 452)* w/ structural features (n= 13).*

| Model | Balanced Accuracy | F1-Score | Precision | Recall | ROC-AUC |
|---|---|---|---|---|---|
| Random Forest | 0.586 | 0.441 | 0.448 | 0.433 | 0.654 |
| XG Boost | 0.575 | 0.420 | 0.451 | 0.403 | 0.641 |
| Gradient Boosting | 0.553 | 0.357 | 0.439 | 0.318 | 0.644 |

**Certain isoforms lack seven transmembrane domains and do not contain a full set of topological features. For this reason, they were excluded from model training. GPCRs we considered to be encoded by stable transcripts if recombinant cells expressing these cells were not statistically enriched within the sub-population of low-GFP cells.*

### Table S2. Relative Contribution of Features in Different Models

| Rank | All GPCRs All Features | | High-GFP GPCRs Structure + Topology | | High-GFP GPCRs Topology Features | | High-GFP GPCRs Structural Features | |
|---|---|---|---|---|---|---|---|---|
| | Feature | Feature Importance | Feature | Feature Importance | Feature | Feature Importance | Feature | Feature Importance |
| 1 | Charged SASA | 0.06885 | $\Delta G_{app,pred}$ TMD6 | 0.080533 | $\Delta G_{app,pred}$ TMD6 | 0.099171 | Total SASA | 0.113689 |
| 2 | Total SASA | 0.063461 | Total SASA | 0.048065 | C-term Loop Length | 0.063678 | isolated beta bridge | 0.092045 |
| 3 | sasa aplr | 0.044372 | N-term loop length | 0.042465 | N-term Loop Length | 0.056413 | turn | 0.091883 |
| 4 | $\Delta G_{app,pred}$ TMD6 | 0.041741 | ECL2 Length | 0.040885 | Molecular weight | 0.050984 | Charged SASA | 0.09081 |
| 5 | coil | 0.037984 | C-term Loop Length | 0.03667 | ECL2 Length | 0.050774 | $3_{10}$ Helix Content | 0.080018 |
| 6 | Molecular weight | 0.033951 | turn | 0.036154 | TMD3 Length | 0.043721 | coil | 0.077969 |
| 7 | ECL2 Length | 0.032032 | $3_{10}$ Helix Content | 0.03605 | TMD5 Length | 0.042167 | sasa aplr | 0.074445 |
| 8 | TMD3 Length | 0.031234 | Molecular weight | 0.034632 | $\Delta G_{app,pred}$ TMD2 | 0.04085 | Alpha Helix | 0.07385 |
| 9 | Alpha Helix | 0.030254 | Charged SASA | 0.033908 | $\Delta G_{app,pred}$ TMD4 | 0.039665 | f_crg | 0.066522 |
| 10 | C-term loop length | 0.028549 | sasa aplr | 0.025969 | TMD1 Length | 0.033791 | sasa_plr | 0.065592 |
| 11 | isolated beta bridge | 0.026309 | instability_index | 0.02442 | Aromaticity | 0.032786 | extended_configuration | 0.06172 |
| 12 | N-term loop length | 0.022469 | $\Delta G_{app,pred}$ TMD2 | 0.023666 | ECL 3 | 0.032244 | f_aplr | 0.056127 |
| 13 | sasa_plr | 0.022323 | Alpha Helix | 0.023285 | gravy | 0.032187 | f_plr | 0.055329 |
| 14 | gravy | 0.021966 | $\Delta G_{app,pred}$ TMD4 | 0.022707 | TMD2 Length | 0.032046 | | |
| 15 | TMD1 Length | 0.021778 | coil | 0.022491 | instability_index | 0.031132 | | |
| 16 | $3_{10}$ Helix Content | 0.018651 | Isoelectric_Point | 0.022379 | Isoelectric_Point | 0.030519 | | |
| 17 | TMD6 Length | 0.016432 | Length TMD5 | 0.021388 | $\Delta G_{app,pred}$ TMD1 | 0.029213 | | |
| 18 | TMD5 Length | 0.016409 | isolated beta bridge | 0.021379 | $\Delta G_{app,pred}$ TMD7 | 0.028785 | | |
| 19 | $\Delta G_{app,pred}$ TMD4 | 0.014686 | TMD3 Length | 0.021124 | TMD6 Length | 0.028595 | | |
| 20 | TMD2 Length | 0.014626 | gravy | 0.02077 | ICL3 Length | 0.02731 | | |
| 21 | ICL1 Length | 0.014592 | Aromaticity | 0.02071 | ECL1 Length | 0.026695 | | |
| 22 | turn | 0.014479 | TMD2 Length | 0.020583 | TMD4 Length | 0.025685 | | |
| 23 | extended_configuration | 0.013478 | ICL2 Length | 0.020281 | ICL2 Length | 0.024597 | | |
| 24 | Aromaticity | 0.01305 | ECL3 Length | 0.02024 | ICL1 Length | 0.024527 | | |
| 25 | minRNAss | 0.012463 | $\Delta G_{app,pred}$ TMD5 | 0.019679 | $\Delta G_{app,pred}$ TMD3 | 0.024473 | | |
| 26 | TMD4 Length | 0.012326 | TMD1 Length | 0.0195 | TMD7 Length | 0.024021 | | |
| 27 | instability_index | 0.012162 | ICL1 Length | 0.019099 | $\Delta G_{app,pred}$ TMD5 | 0.023968 | | |
| 28 | ICL3 Length | 0.012093 | f_crg | 0.018957 | | | | |

| | | | | |
|---|---|---|---|---|
| 29 | $\Delta G_{app,pred}$ TMD2 | 0.011758 | sasa_plr | 0.018542 |
| 30 | $\Delta G_{app,pred}$ TMD7 | 0.010102 | f_aplr | 0.017918 |
| 31 | X40freqens | 0.009905 | $\Delta G_{app,pred}$ TMD3 | 0.017857 |
| 32 | ICL3 Length | 0.009555 | ICL3 Length | 0.017462 |
| 33 | $\Delta G_{app,pred}$ TMD3 | 0.009227 | TMD7 Length | 0.017461 |
| 34 | zeroto38q25RNAss | 0.008998 | TMD6 Length | 0.017086 |
| 35 | ECL1 Length | 0.008905 | extended_configuration | 0.016663 |
| 36 | $\Delta G_{app,pred}$ TMD1 | 0.008561 | $\Delta G_{app,pred}$ TMD1 | 0.016523 |
| 37 | ECL2 Length | 0.008498 | f_plr | 0.015895 |
| 38 | Isoelectric_Point | 0.008326 | $\Delta G_{app,pred}$ TMD7 | 0.01587 |
| 39 | maxRNAss | 0.008219 | TMD4 Length | 0.015527 |
| 40 | zeroto38minRNAss | 0.007767 | ECL1 Length | 0.015207 |
| 41 | X40deltaG | 0.007654 | | |
| 42 | zeroto38maxRNAss | 0.007625 | | |
| 43 | zeroto38avgRNAss | 0.007579 | | |
| 44 | $\Delta G_{app,pred}$ TMD5 | 0.007501 | | |
| 45 | TMD7 Length | 0.006926 | | |
| 46 | q25RNAss | 0.006914 | | |
| 47 | tAI10Min | 0.006903 | | |
| 48 | tAI10q25.25. | 0.006782 | | |
| 49 | plus10valRNAss | 0.006613 | | |
| 50 | CpG_frame1_2 | 0.006416 | | |
| 51 | avgCU_first5 | 0.005919 | | |
| 52 | avgCU_first10 | 0.0059 | | |
| 53 | avgRNAss | 0.005713 | | |
| 54 | GC10min | 0.005621 | | |
| 55 | q75RNAss | 0.005361 | | |
| 56 | tAI10q75.75. | 0.005278 | | |
| 57 | avgCU | 0.005229 | | |
| 58 | Nc | 0.005097 | | |
| 59 | CAI | 0.005036 | | |
| 60 | freqens | 0.005031 | | |
| 61 | zeroto38q75RNAss | 0.004906 | | |

| | | |
|---|---|---|
| 62 | deltaG | 0.00482 |
| 63 | CPSpL | 0.004742 |
| 64 | avgCU_first 20 | 0.004732 |
| 65 | CPS_sum | 0.00466 |
| 66 | GC10max | 0.004538 |
| 67 | GC10q75 | 0.004172 |
| 68 | GC10q25 | 0.00414 |
| 69 | Global_tAI | 0.004113 |
| 70 | GC3s | 0.003927 |
| 71 | tAI10Max | 0.00391 |
| 72 | GC | 0.003705 |

**Table S3. Overview of Machine Learning Features**

| Transcript Feature Name | Description |
|---|---|
| CAI | Codon Adaptation Index. Category: Overall Codon Usage. Calculation Method/Tool: codonW 1.4.4 |
| Nc | Effective number of codons. Category: Overall Codon Usage. Calculation Method/Tool: codonW 1.4.4 |
| GC3s | GC content at the synonymous position. Category: Nucleotide. Calculation Method/Tool: Custom |
| CpG_frame1_2 | Frequency of CG di-nucleotides. Category: Nucleotide. Calculation Method/Tool: Custom |
| avgCU | Average Codon Usage. Category: 5' Codon Usage. Calculation Method/Tool: Biopython |
| CPS_sum | Sum of Codon Pair Score values. Category: Codon Pair Score. Calculation Method/Tool: Coleman et al., 2008 |
| CPSpL | Codon Pair Bias. Category: Codon Pair Score. Calculation Method/Tool: Coleman et al., 2008 |
| Global_tAI | tRNA Adaptation Index. Category: tRNA Adaptation Index. Calculation Method/Tool: codonR |
| tAI10Min | Minimum tAI score over 10 codon windows. Category: tRNA Adaptation Index. Calculation Method/Tool: codonR |
| tAI10Max | Maximum tAI score over 10 codon windows. Category: tRNA Adaptation Index. Calculation Method/Tool: codonR |
| TAI10q25.25. | 25th percentile of tAI scores over 10 codon windows. Category: tRNA Adaptation Index. Calculation Method/Tool: codonR |
| TAI10q75.75. | 75th percentile of tAI scores over 10 codon windows. Category: tRNA Adaptation Index. Calculation Method/Tool: codonR |
| avgCU_first20 | Codon Usage over the first 20 codons. Category: 5' Codon Usage. Calculation Method/Tool: Biopython |
| avgCU_first5 | Codon Usage over the first 5 codons. Category: 5' Codon Usage. Calculation Method/Tool: Biopython |
| avgCU_first10 | Codon Usage over the first 10 codons. Category: 5' Codon Usage. Calculation Method/Tool: Biopython |
| GC | Overall GC content. Category: Nucleotide. Calculation Method/Tool: Custom |
| GC10min | Minimum %GC over 10 codon windows. Category: Nucleotide. Calculation Method/Tool: Custom |
| GC10q25 | 25th percentile of %GC over 10 codon windows. Category: Nucleotide. Calculation Method/Tool: Custom |
| GC10q75 | 75th percentile of %GC over 10 codon windows. Category: Nucleotide. Calculation Method/Tool: Custom |
| GC10max | Maximum %GC over 10 codon windows. Category: Nucleotide. Calculation Method/Tool: Custom |
| X40deltaG | ŒîG of the lowest free energy structure for the first 40 codons. Category: 5' RNA Structure. Calculation Method/Tool: RNAfold |
| X40freqens | Frequency of the lowest free energy structure within the ensemble for the first 40 codons. Category: 5' RNA Structure. Calculation Method/Tool: RNAfold |
| plus10valRNAss | Average hybridization probability centered around +10 base, i.e. average of +5 to +15. Category: 5' RNA Structure. Calculation Method/Tool: RNAfold |
| zeroto38avgRNAss | Average hybridization probability over 10 base windows from 0 to +38. Category: 5' RNA Structure. Calculation Method/Tool: NUPACK |

| | |
|---|---|
| zeroto38minRNAss | Minimum hybridization probability over 10 base windows from 0 to +38. Category: 5' RNA Structure. Calculation Method/Tool: NUPACK |
| zeroto38q25RNAss | 25th percentile of hybridization probability over 10 base windows from 0 to +38. Category: 5' RNA Structure. Calculation Method/Tool: NUPACK |
| zeroto38q75RNAss | 75th percentile of hybridization probability over 10 base windows from 0 to +38. Category: 5' RNA Structure. Calculation Method/Tool: NUPACK |
| zeroto38maxRNAss | Maximum hybridization probability over 10 base windows from 0 to +38. Category: 5' RNA Structure. Calculation Method/Tool: NUPACK |
| deltaG | ŒîG of the lowest free energy structure. Category: Overall RNA Structure. Calculation Method/Tool: RNAfold |
| freqens | Frequency of the lowest free energy structure within the ensemble. Category: Overall RNA Structure. Calculation Method/Tool: RNAfold |
| avgRNAss | Average hybridization probability over 10 base windows. Category: Overall RNA Structure. Calculation Method/Tool: NUPACK |
| minRNAss | Minimum hybridization probability over 10 base windows. Category: Overall RNA Structure. Calculation Method/Tool: NUPACK |
| q25RNAss | 25th percentile of hybridization probability over 10 base windows. Category: Overall RNA Structure. Calculation Method/Tool: NUPACK |
| q75RNAss | 75th percentile of hybridization probability over 10 base windows. Category: Overall RNA Structure. Calculation Method/Tool: NUPACK |
| maxRNAss | Maximum hybridization probability over 10 base windows. Category: Overall RNA Structure. Calculation Method/Tool: NUPACK |

| Protein Feature name | Description |
|---|---|
| Length_without_SP | Length of the protein after removal of the signal peptide. |
| TMD1-7 Length | Normalized lengths of each transmembrane domain (TMD1–TMD7), relative to the full-length protein. TMDs are predicted using **DeepTMHMM**, and their coordinates are refined using the **von Heijne transfer free energy scale** von_heijne_scan_functions_v39_signal_pep_topcons.py (https://github.com/schebachlab/PRF-Search).<br>The lengths are then normalized to the length of protein |
| N-term Loop Length | Normalized length of the N-terminal extracellular region preceding TMD1. |
| ICL1-3 Length | Normalized lengths of the intracellular loops immediately following TMD1, TMD2, and TMD3, respectively. |
| ECL1-3 Length | Normalized lengths of the extracellular loops immediately preceding TMD2, TMD3, and TMD4, respectively. |
| C-term Loop Length | Normalized length of the C-terminal intracellular region following TMD7. |
| $\Delta G_{app,pred}$ TMD1-7 | Free energy of membrane insertion (ΔGAPP,PRED) for each TMD (TMD1–TMD7), computed using the **von Heijne transfer free energy scale** via von_heijne_scan_functions_v39_signal_pep_topcons.py (https://github.com/schebachlab/PRF-Search). These ΔGAPP,PRED values reflect the biophysical favorability of each TMD's insertion into the membrane. |
| Molecular_Weight | The molecular weight of the protein (in Daltons), computed from its amino acid composition. |
| Isoelectric_Point | The theoretical pH at which the protein carries no net charge. |
| Aromaticity | The fraction of aromatic amino acids (Phe, Trp, Tyr) present in the protein. |
| instability_index | Predicts the in vitro stability of a protein based on dipeptide composition. Values greater<br>than 40 suggest stability; values less 40 indicate instability. Defined by Guruprasad et<br>al. (1990):(https://pubmed.ncbi.nlm.nih.gov/2075190/) |
| gravy | Average hydrophobicity of the amino acids in the sequence using the Kyte–Doolittle<br>scale: https://doi.org/10.1016/0022-2836(82)90515-0. |

| | |
|---|---|
| Total SASA | The total solvent-accessible surface area (SASA) of the protein, calculated using FreeSASA:<br>https://freesasa.github.io/. |
| sasa_crg<br>sasa_plr<br>sasa_aplr | Contributions to total SASA from:<br>– Charged SASA: Charged residues (Asp, Glu, Lys, Arg,His).<br>– SASA plr: Polar residues (Ser, Thr, Asn, Gln, Tyr, Cys).<br>– SASA aplr: Apolar (non-polar) residues (Ala, Val, Ile, Leu, Met, Phe, Trp, Pro,<br>Gly). |
| f_crg<br>f_plr<br>f_aplr | Fractions of the total SASA contributed by charged, polar, and apolar residues, respectively<br>(range: 0–1). |
| alpha_helix | Fraction of residues in alpha-helical conformation, as assigned by STRIDE. |
| $3_{10}$ Helix Content | Fraction of residues in $3_{10}$ helix conformation. |
| extended_configuration | Fraction of residues in extended strand (beta-sheet-like) conformations. |
| isolated_beta_bridge | Fraction of residues forming isolated beta-bridges (not part of extended beta-sheets). |
| turn | Fraction of residues in turns—tight loops reversing the direction of the peptide backbone. |
| coil | Fraction of residues in unordered, coil-like conformations. |
